# A structure-guided pipeline to uncover the underexplored beta-lactamases from Antarctic and Subantarctic soil microbiota

**DOI:** 10.1101/2025.09.30.679653

**Authors:** José Coche-Miranda, Patricio Arros, Nicolás Canales, Camilo Berríos-Pastén, Rosalba Lagos, Julieta Orlando, Francisco P. Chávez, Andrés E. Marcoleta

## Abstract

As the One Health approach points out, the environment can significantly affect human health, particularly having diverse and profound implications in the ongoing antimicrobial resistance crisis. One central aspect is the potential role of environmental microorganisms as a source of resistance genes that could emerge among pathogens aided by horizontal transfer. In this context, previous reports showed that the Antarctic soil microbiota hosts a rich resistome, including putative beta-lactamases conferring resistance to beta-lactams, the most widely used antibiotics for treating bacterial infections worldwide. However, the diversity of beta-lactamases across different areas of Antarctica and Subantarctic islands and their associated microbiota remains largely unexplored. In this study, we analyzed an extensive collection of Antarctic soil metagenomes, applying a novel bioinformatic pipeline based on combining sequence identity and structural similarity criteria to search for *bona fide* distant beta-lactamase orthologs. We found several classes and families, with a notable predominance of class-B metallo-beta-lactamases, including proposed novel families and variants of known families. The beta-lactamase diversity and dominant classes varied across sites following changes in the microbial community, observing three main sample groups: 1) Subantarctic islands, 2) Antarctic Peninsula and surrounding islands; and 3) Cold-desert environments. The most prevalent beta-lactamases corresponded to subclass B3. We reconstructed more than 3500 metagenome-assembled genomes (MAGs) and searched for beta-lactamases among them. The main taxa encoding the beta-lactamases were Pseudomonadota and Bacteroidota. Moreover, a certain proportion of beta-lactamases was associated with mobile genetic elements, especially integrons and insertion sequences, suggesting their potential dissemination by horizontal transference. This evidence reinforces the role of the environment, and especially (sub)Antarctic soils, as a reservoir of resistance genes, in particular, beta-lactamases.

## INTRODUCTION

We are currently facing the global antimicrobial resistance (AMR) crisis, where, as estimated by the Global Burden of Disease (GBD) 2021 Antimicrobial Resistance Collaborators, about 1.3 million people die each year due to AMR-associated infections. This number could reach 10 million people by 2050 if adequate preventive measures are not taken (Naghavi et al., 2024), highlighting the importance of understanding the origin and dynamics of AMR. Given that AMR is a complex multidimensional issue, it should be faced following the “One Health” paradigm that considers the interrelationship between the environment, animals, and humans (Larsson et al., 2023). Within this context, it has been widely demonstrated that the antibiotic resistance genes are ancient and ubiquitous, and that the microbiota living in the wider environment, even in remote, pristine, and extreme areas, can act as a source of ARGs that bacterial pathogens could eventually acquire (Agudo and Reche, 2024; D’Costa et al., 2011).

Among the ARGs of greatest concern are those encoding beta-lactamase (BLA) enzymes, the most widespread and efficient mechanism within Gram-negative bacteria to resist beta-lactam antibiotics (de Souza et al., 2024; Shahid et al., 2022; Zeng and Lin, 2013), which correspond to the most commonly used antimicrobial therapy in clinical settings (Mora-Ochomogo and Lohans, 2021). BLAs are grouped into two major categories based on sequence similarity and structural characteristics: serine beta-lactamases (SBLs), characterized by sharing a serine residue essential for their mechanism of action, including classes A, C, and D; and metallo-beta-lactamases (MBLs), which utilize zinc ions to catalyze the hydrolysis of beta-lactam antibiotics, encompassing subclasses B1, B2, and B3 (Akhtar et al., 2022; Bush and Jacoby, 2010).

Like other resistance mechanisms, BLAs are ubiquitous and ancestral, existing for approximately 2 billion years in the case of SBLs and 2.5 billion years for MBLs (Bush, 2018; Fröhlich et al., 2021). While their primary role in evolution remains unknown, it is important to note that both groups belong to protein superfamilies with a wide range of functions. SBLs belong to the PBP-like superfamily, characterized by alpha-beta-alpha structural fold and conserved SXXK and semi-conserved SXN and KTG(T/S) active site motifs (Zapun et al., 2008). In contrast, MBLs belong to the MBL-fold-like superfamily, characterized by an alpha-beta-beta-alpha-fold, and two conserved active-site metal-binding motifs involving H116/H118/H196 and D120/H121/H263, along with D221 that coordinates both metals (Fröhlich et al., 2021).

BLA genes have been identified in various environments with or without human intervention, including remote and extreme areas (Allen et al., 2009; Bhullar et al., 2012). Among them, soils from maritime Antarctica have been described as a rich source of potential ARGs, including BLAs, correlating with the presence of bacteria resistant to beta-lactams and several other antibiotics (Goethem et al., 2018; Marcoleta et al., 2022; Santos et al., 2022). However, their mechanisms have not been studied in depth, and the selective pressure driving this resistance remains unclear.

A beta-lactamase survey in eleven metagenomes from Antarctic microbial mats along a latitudinal gradient found putative BLA genes from all four classes (A–D), with class C being the most abundant (Azziz et al., 2019). However, the distribution and abundance of potential BLAs in soils from different areas of Antarctica, which would have higher microbial and gene diversity compared to microbial mats, remain unknown. Furthermore, this study did not address which microbial taxa harbor these BLA genes, obscuring their evolutionary trajectories and ecological roles. Moreover, although the divergence of environmental β-lactamase sequences from reference clusters suggests evolutionary novelty, their structural folds, active site motifs, and phylogenetic placement remain to be clarified.

Identifying β-lactamase orthologs solely through amino acid sequence identity presents a notable limitation, particularly when investigating deeply divergent or environmentally embedded variants. Many remote or ancestral β-lactamase genes exhibit low sequence similarity to clinically characterized enzymes, falling below detection thresholds used by standard annotation pipelines. As a result, these cryptic orthologs remain undetected or misclassified. This obstacle can be subverted by leveraging structure prediction algorithms such as AlphaFold or ESMFold. By modeling protein tertiary structures and identifying conserved structural motifs, it is possible to unveil functional homologs hidden within environmental metagenomes, even in cases where sequence identity drops below 25% (Fröhlich et al., 2021). In this line, we hypothesized that a large variety of beta-lactamases, including potentially novel families, are encoded by Antarctic soil bacteria, which can be surveyed using a tailored bioinformatic pipeline to find remote environmental beta-lactamase orthologs from metagenomic data.

In this work, we devised a bioinformatic workflow to screen for *bona fide* beta-lactamase orthologs, applying it to an extensive collection of Antarctic soil metagenomes, covering sub-Antarctic islands, maritime Antarctica, and high-latitude cold desert environments. We assessed the diversity, abundance, and geographical distribution of 1916 putative beta-lactamases (classes A, B1, B2, B3, C, and D) from 81 metagenomes of soil samples collected from Antarctica. These results provide relevant insights into the potential of Antarctic soil microbiota as a reservoir of beta-lactamases that could eventually emerge among pathogens, aggravating the ongoing AMR crisis.

## RESULTS

### Towards a metagenomic survey of Antarctic soil microbiota and their beta-lactamases

We built a set of 81 Antarctic and Subantarctic soil metagenomes, corresponding to most of the available until September 2024 in the NCBI database, plus 12 metagenomes sequenced by our group. We excluded those who did not meet the quality criteria (see Methods) or lacking GPS coordinates of the respective sampled point. These included six metagenomes from sub-Antarctic Islands (SI), namely South Georgia, Marion, Kerguelen, Bartolome, Possession, and Bouvetoya islands. Also, 37 from the Antarctic Peninsula and surrounding islands (AP), mainly the South Shetland Archipelago, and 38 samples from high-latitude cold desert areas (CD), mainly the McMurdo Dry Valleys, the Shackleton Glacier, and the Union Glacier (Figure 1A). The complete list of metagenomes included in this study, along with their accession numbers and relevant metadata, is provided in Table S1.

**Figure 1.**
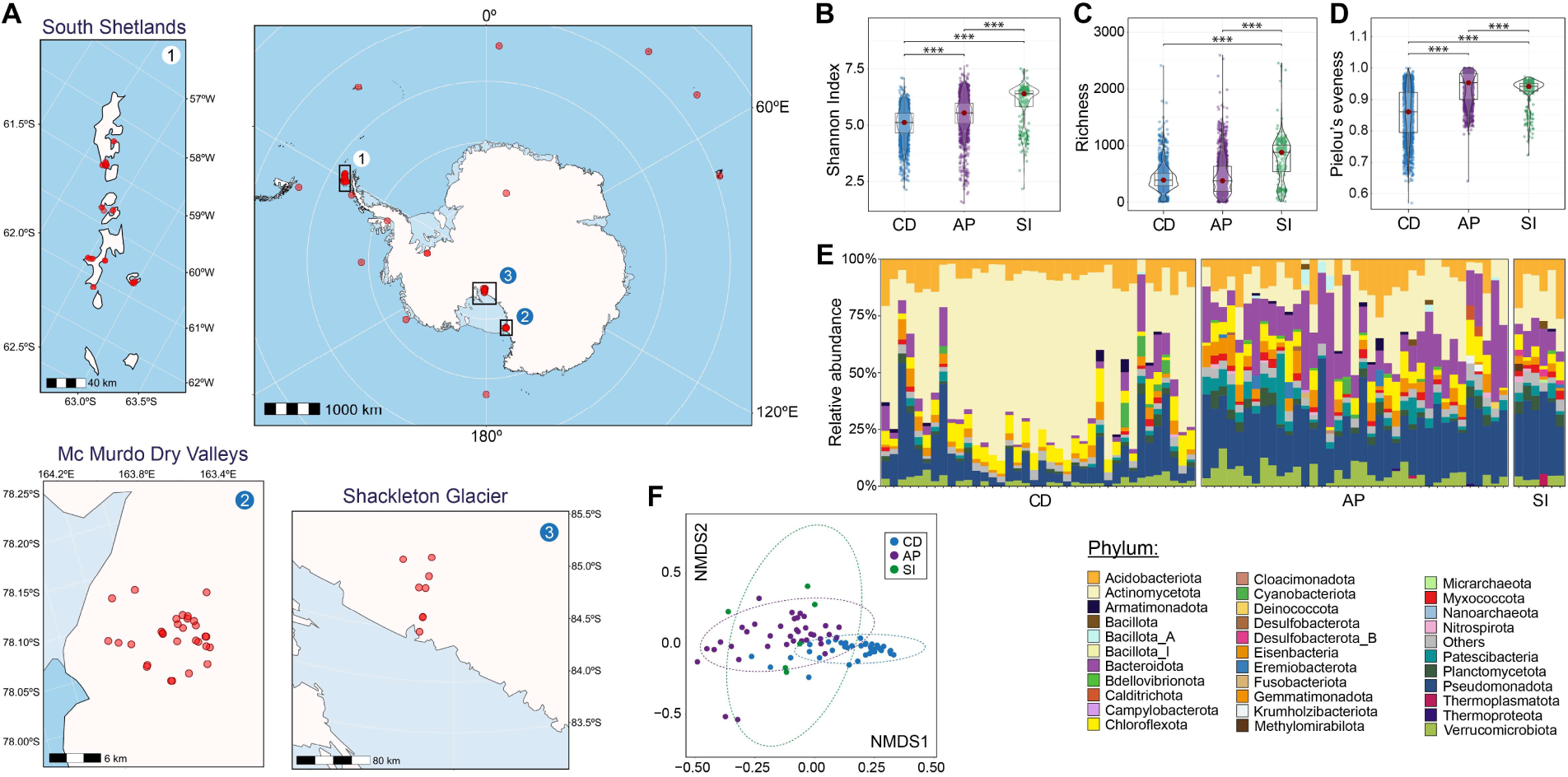
Geographic location and microbial diversity of the soil metagenomes included in this study. A: Map showing the location of the soils represented in our metagenome set. The insets (1-3) show a zoomed view of the areas concentrating a larger number of samples. B, C, D: distribution of diversity indices across cold desert (CD), Antarctic Peninsula and surrounding islands (AP), and subantarctic islands (SI) soils, based on 43 essential single-copy marker genes. \*\*\**p*_FDR-adjusted_<0.001 (Dunn’s pairwise test). E: Relative abundance of microbial taxa at the phylum level. F: Non-Metric Multidimensional Scaling based on the Bray-Curtis beta-dissimilarity at the family level across the Antarctic and subantarctic soils. Significant differences were observed between AP vs. CD (\*\*\**p*_FDR-adjusted_=0.000999) and CD vs. SI (\*\*\**p*_FDR-adjusted_=0.000999), as determined by Pairwiseadonis.

First, we aimed to explore the diversity and taxonomic composition of the microbial communities inhabiting the different Antarctic soils studied. For this, we used the SingleM pipeline, specially designed for microbial diversity analyses from shotgun metagenomic data based on a collection of 59 essential single-copy genes (ESCGs) (Woodcroft et al., 2025). However, we only kept the results for 43 ESCGs due to 16 not being observed in at least one sample. Shannon Index values indicated the least total diversity in CD soils (median: 5.12), being significantly higher in AP soils (median: 5.55), and in SI soils (median: 6.40) (Figure 1B).

This observation could be explained by a similar richness in CD and AP soils (median: 393 and 381, respectively), which is significantly higher in SI soils (median: 882) (Figure 1C), together with a high evenness in AP and SI (median: 0.95 and 0.94, respectively), being significantly lower in CD (Figure 1D).

Regarding the microbial community structure, the SI and AP soils showed a more similar phyla composition, predominating Pseudomonadota (mainly the order Burkholderiales), Bacteroidota (mainly Chitinophagales), Acidobacteriota (mainly Pyrimonadales), and, to a lesser extent, Actinomycetota (mainly Acidimicrobiales and Pyrinomondales) (Figure 1E). Conversely, in most CD soils, Actinomycetota (mainly the order Solirubrobacterales) largely predominated, with a markedly lower abundance of Bacteroidota, Pseudomonadota, Gemmatimonadota, and Patescibacteria. In agreement, beta-diversity dissimilarity analysis at the family level showed a more similar taxonomic composition between SI and AP soils, while the CD soils clustered apart, given their different community structure (Figure 1F).

### Identification of putative beta-lactamases in Antarctic and Subantarctic soil metagenomes: a tailored pipeline for remote ortholog prediction

A critical obstacle to searching for orthologs of defined proteins (e.g., antimicrobial resistance factors) in metagenomes from remote and poorly studied environments, such as Antarctica, is the low sequence similarity of the environmental proteome with proteins present in public databases (mostly from human-associated sources). Therefore, protein searches based solely on amino acid sequence comparison often lead to high false positive rates, given the restriction of working in the “twilight zone” of sequence identity. This caveat is especially relevant for protein families harboring diverse functions, such as PBP-like and MBL-fold to which beta-lactamases belong, where enzymes sharing a certain degree of sequence similarity can have markedly different activities. To overcome these limitations, we devised a bioinformatic workflow named PROSSAF (PRedicting Orthologs by combining Sequence, Structure, and Annotation of Function) (Figure 2). This pipeline builds and integrates sequence, structure, and functional similarity embeddings to generate confident protein ortholog clusters, aiming to distinguish true orthologs from related proteins having different functions. Moreover, this pipeline combines different protein assembly and clustering strategies to maximize the proteins identified from metagenomic data. We applied this pipeline to thoroughly search for putative beta-lactamases among the Antarctic and subantarctic soil metagenomes.

**Figure 2.**
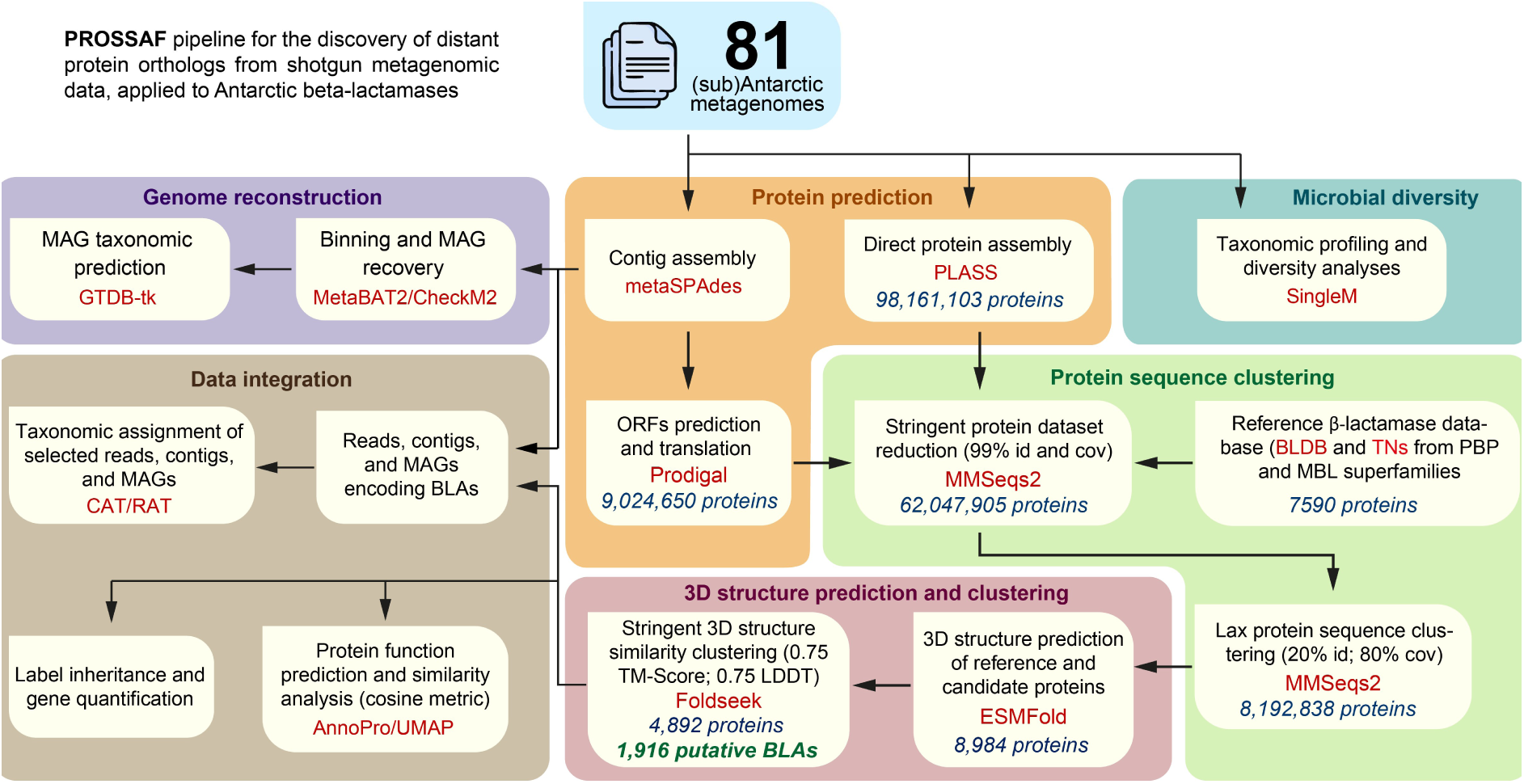
PROSSAF workflow for metagenomic Antarctic beta-lactamase survey and microbial diversity analysis. The main stages and aims of the workflow are shaded in colors. The main bioinformatic tools used for the different stages are indicated in red. The numbers of proteins obtained in part of the steps are indicated in blue.

Based on the reads from the 81 metagenomes, first, a contig-level assembly was performed with metaSPAdes, followed by the prediction of ORFs in the contigs, resulting in a total of approximately 9 million proteins. Also, direct protein assemblies from the reads were carried out using the PLASS program (Steinegger et al., 2019). PLASS enables ultra-fast, scalable protein-level assembly, overcoming the biases in contig reconstruction to uncover low-abundance proteins, protein variants, and novel sequences often missed by traditional workflows. This way, roughly 98 million proteins were predicted, increasing the predicted proteins at the contig level by tenfold.

To identify putative BLAs, the Beta-Lactamase DataBase (BLDB, http://bldb.eu/) was used as a reference (7566 sequences clustered into 5458 non-redundant representatives). Additionally, as negative controls, we used a set of proteins phylogenetically related to BLAs from the PBP-like and MBL-fold superfamilies but demonstrated to have different functions (6 and 18 sequences, respectively) (“true negatives”, Table S2). To eliminate sequence redundancy, all the metagenomic and reference proteins were clustered at 99% sequence identity and 99% bidirectional coverage, resulting in a total of approximately 62 million non-redundant proteins. To identify an initial set of Antarctic beta-lactamases, the previous set, including the positive and negative reference proteins, was clustered at 20% sequence identity and 80% bidirectional coverage, resulting in around 8 million proteins, of which 3502 clustered either with reference BLAs (2181 sequences) or with true negatives (1321 sequences).

Finally, the three-dimensional structures of all 3502 Antarctic proteins, 5428 non-redundant reference BLAs and 24 TNs were predicted, and structure-guided clustering was performed using strict local and global similarity criteria (TM-score ≥ 0.75, lDDT ≥ 0.75). This step restricted twilight-zone sequence alignments with complementary steric constraints imposed by known structural motifs observed in the reference BLAs and TNs. As a result, we obtained a final set of 1916 predicted non-redundant Antarctic beta-lactamase orthologs forming 25 distinct clusters with reference proteins (Figure 2; Table I), and 692 Antarctic TN orthologs forming 9 clusters. No mixed clusters containing both reference BLAs and TNs were observed, demonstrating the robustness of the developed PROSSAF pipeline.

The 1916 putative BLAs, according to sequence and structural criteria, were grouped into 986 serine beta-lactamases (SBLs) and 930 metallo-beta-lactamases (MBLs). Among the SBLs, 512 sequences were categorized as Class A, 77 as Class C, and 397 as Class D, with sequence identity ranging from 41.1% to 100% compared to BLDB proteins. Six Class A sequences shared 97% to 100% identity with BLDB proteins (five TEM variants and CARB-16), while in Class D, a single sequence with 99.6% identity to OXA-455 was identified. These sequences are likely to correspond mainly to BLAs disseminated from other environments, as exemplified by the TEM-116 variant identified in this study, which has been reported both in clinical settings (Jeong et al., 2004; Usha et al., 2008) and in non-clinical environments (Maravić et al., 2016; Mondal et al., 2019; Naidoo et al., 2020), attributed to its plasmid-mediated mobilization. For MBLs, 263 sequences were classified into subclass B1, two into subclass B2, and 665 into subclass B3, with sequence identity ranging from 41.3% to 89.5% compared to beta-lactamases from BLDB (Table 1).

**Table 1.** Predicted beta-lactamase orthologs in the Antarctic and Subantarctic soil metagenomes.

| Beta-lactamase category | Structural class | N° of sequences | N° of sequences ($\geq 90\%$ id) | N° of structural clusters | % sequence identity against BLDB |
| --- | --- | --- | --- | --- | --- |
| SBLs | A | 512 | 128 | 10 | 42.1-100 |
|  | C | 77 | 12 | 1 | 51.1-56.3 |
|  | D | 397 | 72 | 3 | 44.5-99.6 |
|  | **Total** | 986 | 211 | 13 | - |
| MBLs | B1 | 263 | 53 | 3 | 47.7-89.5 |
|  | B2 | 2 | 1 | 1 | 61-62 |
|  | B3 | 665 | 180 | 7 | 41.3-84.8 |
|  | **Total** | 930 | 233 | 11 | - |

Regarding the distribution of non-beta-lactamase proteins from the PBP superfamily, 186 potential orthologs of carboxyl-esterases were found (40 EstU1-like, 138 PBP-5-like, and 8 PBP-A-like). For the MBL-fold superfamily proteins, 665 orthologs were identified, of which 174 could correspond to type II glyoxylases (13 1xm8-like and 161 glob-like), 7 to methyl-parathion hydrolases, 33 to PQQ-biosynthesis protein B, 4 to pyridoxolactonases, and 447 to ribonuclease Z-like proteins (Table S3). Therefore, the PROSSAF pipeline would successfully distinguish between bona fide remote BLA orthologs and closely related proteins with different functions.

To get additional evidence of the potential BLA function of the predicted orthologs, even in the lower sequence identity range, a conserved catalytic motif analysis was performed based on the known molecular features of each structural class. Forty-six out of 51 sequences of class A (40-50% amino acid identity range) possessed the S(70)XXK, S(130)XN, E(166)XXLN, and K(234)TG active site motifs (Figure S1). The differences were observed in the catalytic motif K(234)TG, where three sequences displayed the motif RIG, while two sequences exhibited the motif NTG. Also, the six class C sequences (50-60% amino acid identity) comprised the S(64)XXK, Y(150)XN, and K(315)T/SG active site motifs (Figure S2). Furthermore, 49 out of 55 sequences of class D (44-50% amino acid identity) exhibited the S(70)TFK, S(118)XV, Y/F(144)GN, and K/S(216)TG active site motifs (Figure S3). Differences were mainly identified at valine 120 within the S(118)XV motif, with five sequences showing leucine at position 120 and one sequence showing isoleucine.

In the case of MBLs, 22 out of 23 subclass B1 sequences (47-50% amino acid identity) comprised the H(116)-H(118)-H(196) and D(120)-C(221)-H(263) active site motifs (Figure S4), with one sequence having a Tyr instead of a His residue at position 196. Additionally, 26 out of 30 sequences of subclass B3 (40-50% amino acid identity range) contained the H(116)-H(118)-H(196) and D(120)-H(121)-H(263) active site motifs (Figure S5), where four sequences exhibit a non-conserved substitution at serine 221 (Arg, Met, Gly, or Cys). Finally, in the case of subclass B2, only two sequences were identified in this study, both of which contained the D(120)-C(221)-H(263) active site motif. Therefore, most of the identified BLAs, including those with low sequence identity to reference proteins, exhibit the key motifs typical for the respective structural class. For the remaining sequences, at the highest sequence identity thresholds defined for each class, at least 91% conservation was observed across all of the aforementioned catalytic residues (Table S2).

True protein orthologs are expected to share, to some extent, sequence, structure, and function characteristics. In this line, we performed a multidimensional function prediction on the putative Antarctic BLAs, examining the functional coherence across the ortholog clusters based on sequence and structure similarity. For this, we used the recently developed AnnoPRO pipeline (Zheng et al., 2024) based on multi-scale protein function representation aided by deep learning. In this analysis, each protein is represented by a 6109-dimensional probability vector corresponding to the Gene Ontology (GO) terms from three aspects: molecular function, cellular compartment, and biological process. Then, the similarity among the function vectors from all the 1916 putative BLAs, 692 putative TNs and 2284 reference proteins was calculated and compared using the cosine metric (Koehler et al., 2023), followed by a Uniform Manifold Approximation and Projection for Dimension Reduction (UMAP) plotted in a 2D space (Figure S6).

SBLs clustered separately from all other protein families, with distinct separation between classes A, C, and D. They appeared functionally divergent from both PBP-like and MBL-fold-like true negatives—except for an EstU1-like carboxylesterase, which showed functional similarity to class C SBLs. Carboxylesterases PBP-5-like and PBP-A-like also formed distinct clusters, showing that the complementary function space can distinguish SBLs from other PBP-like and MBL-fold-like proteins independently of the sequence and structure similarities used to identify the Antarctic orthologs. In contrast, most of the MBL-fold-like superfamily of proteins cannot be adequately identified just by functional prediction alone, except for Ribonuclease-Z-like orthologs which form a distinct functional similarity group, showing that the combination of sequence, structure and functional comparisons within the MBL-fold-like superfamily of proteins is necessary to properly distinguish between different orthologous groups, as proposed by the implementation of the complete PROSSAF pipeline.

### Phylogenetic and structural landscape of Antarctic beta-lactamases

We further investigated the main features of the Antarctic BLAs and their phylogenetic relationships with reference beta-lactamases and other enzymes from the PBP-like and MBL-fold superfamilies (true negatives). For this purpose, Antarctic BLAs, together with their closest reference sequences (belonging to the same structural cluster), were clustered at 90% sequence identity and bidirectional coverage. A similar approach was applied to Antarctic proteins that structurally clustered with true negatives. This strategy enabled the analysis of 128 representative Antarctic BLAs belonging to class A, 53 B1, 1 B2, 179 B3, 11 C, and 72 class D, along with 53 Antarctic PBP-like proteins and 116 MBL-fold-like proteins that are not β-lactamases, as well as 587 beta-lactamases representative sequences from BLDB and 14 true negative sequences. Phylogenetic trees were constructed starting from multiple sequence alignments of these *bona fide* Antarctic beta-lactamase orthologs, plus BLDB reference sequences and PBP-like and MBL-fold true negatives. Also, the structural cluster information from the Foldseek-based clustering step in the PROSSAF pipeline was incorporated into the trees.

Regarding the PBP-like superfamily, as expected, class A, class C, and class D reference beta-lactamase sequences formed deeply separated monophyletic clades, each including also Antarctic orthologs (Figure 3). The highest diversity of sequences was observed for class A, followed by class D and then class C. Accordingly, markedly more structural clusters were found for class A (10), only 3 for class D, and one for class C. As demonstrated previously, at the sequence and structural levels, A and C are close to each other while D is more distant (Hall and Barlow, 2004). Conversely, all the non-beta-lactamase PBP-like proteins, except for BlaR1 (class D), formed separate clades from beta-lactamases. Regarding BlaR1, this is a result consistent with previous phylogenetic analyses within the PBP-like superfamily, showing that this protein could represent a paralog that retained structural similarity to class D beta-lactamases but lost the hydrolytic activity, evolving to act as a beta-lactam sensor and signal transducer (Fröhlich et al., 2021).

**Figure 3.**
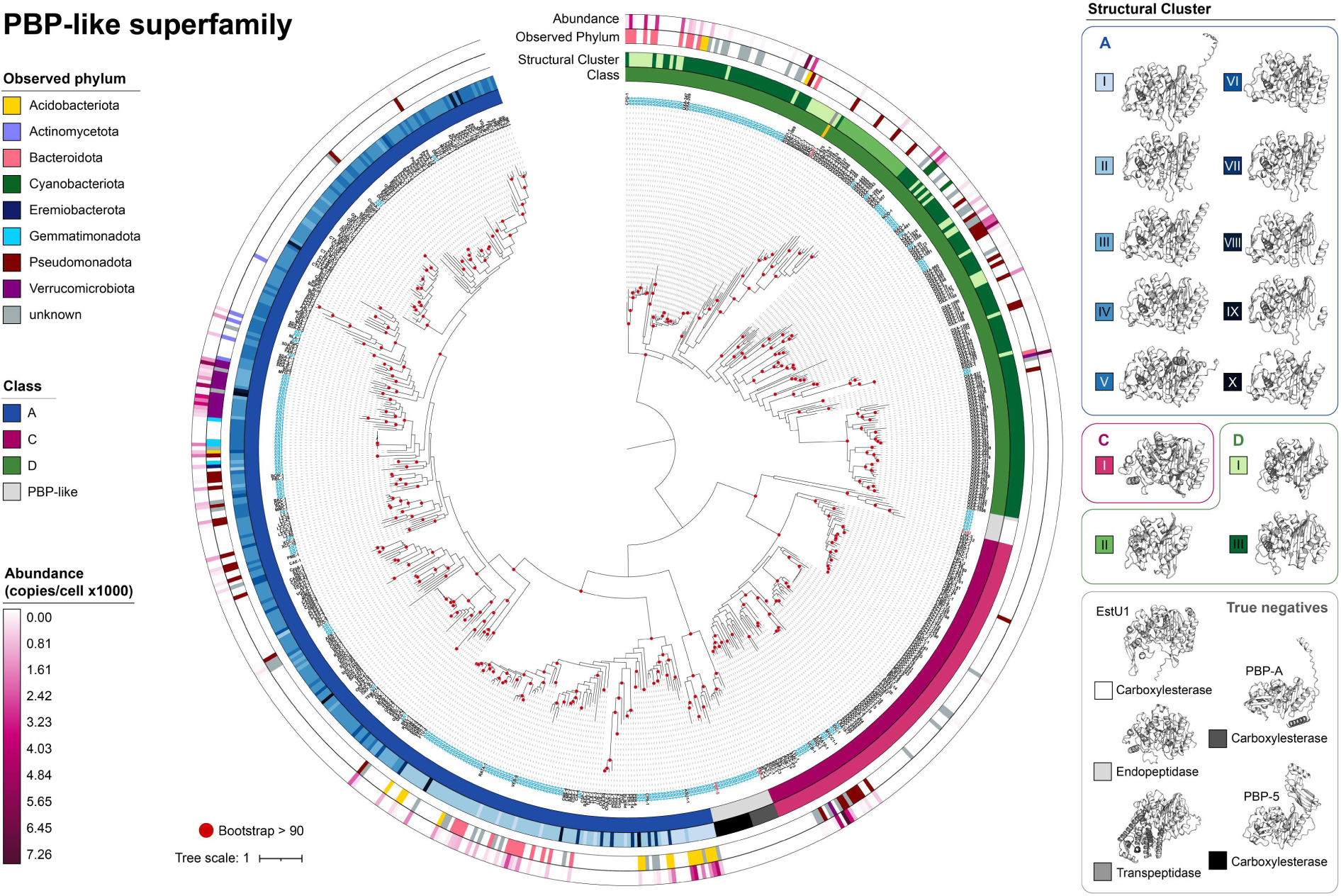
Maximum-likelihood tree showing the phylogenetic relationships, structural features, and relative abundance of representative Antarctic class A, C, and D beta-lactamases and other PBP-like superfamily proteins. Light blue tip labels indicate Antarctic proteins. Red tip labels indicate non-beta-lactamase proteins. Structural clusters were based on a 0.75 LDDT and 0.75 TM-score cutoff. A representative structure from each cluster is shown. The beta-lactamase gene abundances were calculated based on metagenomic reads mapping for each metagenome. The total abundance shown for each protein corresponds to the sum across all the sites.

Inside class A, two major clades were observed. A more numerous one, including most of the references plus some Antarctic, and a smaller one, including a higher proportion of Antarctic sequences. This separation is supported by structural clustering, where the minor clade included primarily sequences from the structural clusters I, II, and III, while, in the major clade, clusters IV, V, and VI predominated. In class C, only one structural cluster was observed, supporting the higher homogeneity inside this group. In class D, three structural clusters were observed relatively intermingled in the tree, all of them with Antarctic orthologs.

For all the classes, it was observed that some Antarctic sequences grouped into clades distinct from BLDB references, which could represent novel beta-lactamase families (Figure 3). Class A beta-lactamases show that most monophyletic clades consist exclusively of Antarctic sequences, sharing 41.1% to 59.8% sequence identity with their closest references. In contrast, clades that share a common ancestor with known references mostly display <70% identity, except for one clade of four sequences related to RATA-1, showing 80.92%–86.51% identity. RATA-1 is a class A beta-lactamase identified from *Riemerella anatipestifer* isolates, with most of its known variants reported in marine *Flavobacteriaceae* (phylum Bacteroidota) (Luo et al., 2024). This finding suggests its presence also in soil-associated environmental bacteria of the same phylum.

Concerning class C BLAs, the monophyletic clades mainly group with known references such as LRA18-1, IDC-1, MYCC1-1, LRA10-1, and UCB-1. Notably, two Antarctic sequences share at least 83% identity with IDC-1, an integron-derived cephalosporinase detected by functional metagenomics of DNA from river sediments contaminated with municipal and hospital sewage (Böhm et al., 2020). Within class D, the largest monophyletic clade, composed solely of Antarctic sequences, comprises 30 BLAs with 48.3%–55.1% identity to the closest references, potentially constituting a novel family. Another clade of four Antarctic sequences clustered with OXA-1091 (70%–74% sequence identity), a beta-lactamase detected in the non-clinical bacterium *Bradyrhizobium* sp. BTAi, which exhibits activity against oxacillin and imipenem (Lupo et al., 2022).

Concerning the MBL-fold superfamily, it was observed that all Antarctic sequences and BLDB reference sequences from subclasses B1, B2, and B3 clustered into separate clades compared to their non-beta-lactamase homologs within the superfamily (Figure 4). Subclass B1 beta-lactamases included a large monophyletic clade of 19 Antarctic sequences sharing 48.5%–58% identity with their closest references, corresponding to a putative novel family. A distinct clade of eight sequences is closely related to MYO-1 (70%–88% identity), a beta-lactamase identified in *Myroides odoratimimus* able to hydrolyze imipenem (Berglund et al., 2017). Additionally, novel Antarctic variants of CHM-1 and CEMC19-1 BLAs, and a highly divergent Antarctic BLA sharing a common ancestor with the CAM-1 carbapenemase found in *P. aeruginosa* (Boyd et al., 2019).

**Figure 4.**
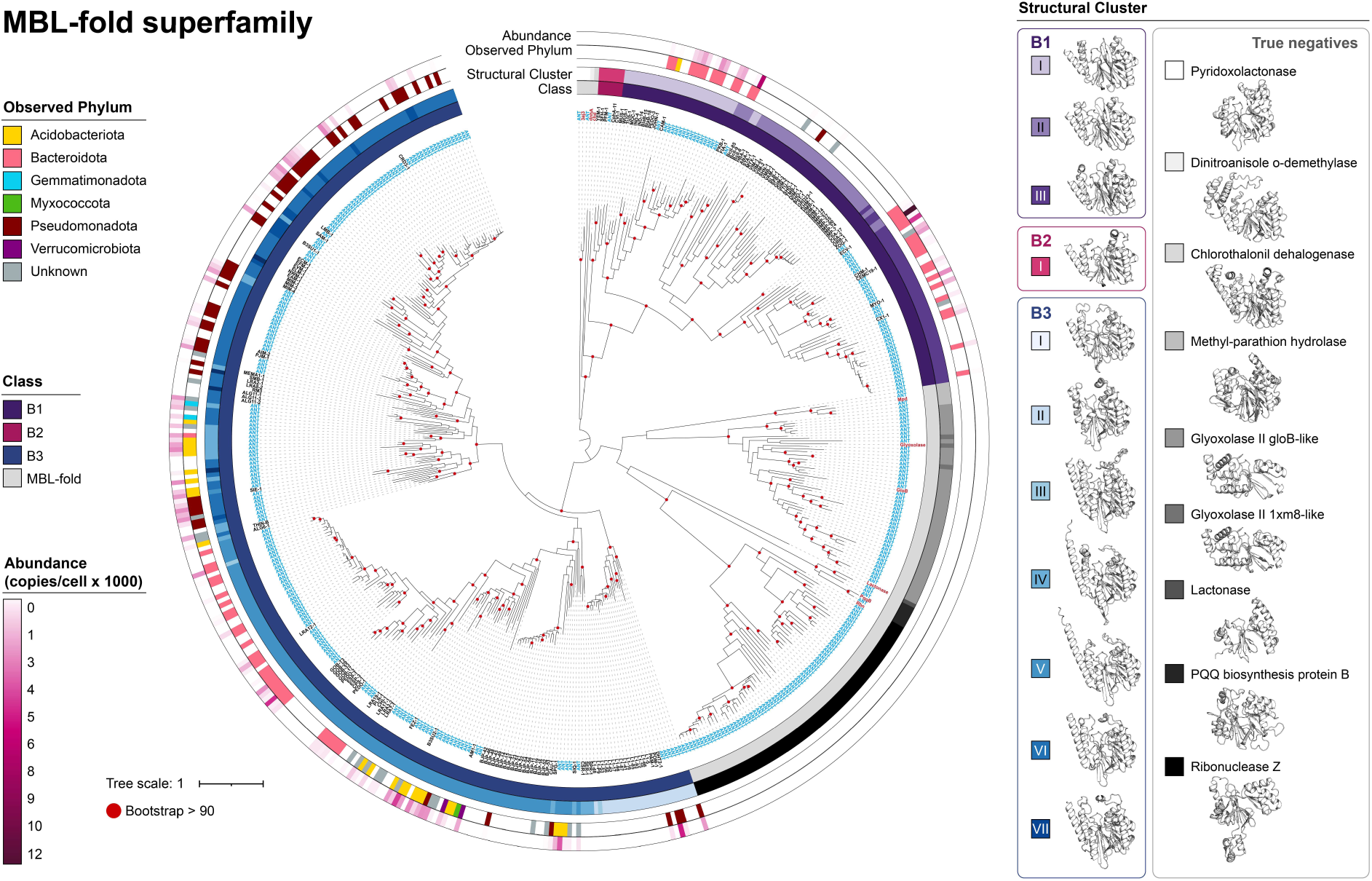
Maximum-likelihood tree showing the phylogenetic relationships, structural features, and relative abundance of representative Antarctic class B beta-lactamases and other MBL-fold superfamily proteins. Light blue tip labels indicate Antarctic proteins. Red tip labels indicate non-beta-lactamase proteins. Structural clusters were based on a 0.75 LDDT and 0.75 TM-score cutoff. A representative structure from each cluster is shown. The beta-lactamase gene abundances were calculated based on metagenomic reads mapping for each metagenome. The total abundance shown for each protein corresponds to the sum across all the sites.

For subclass B3, the largest monophyletic clade contains 28 Antarctic BLAs showing 63.4%–69.4% identity with their nearest reference (LRA12-1) and belonging to structural cluster IV. Remarkably, LRA12-1 was identified through functional metagenomics from a remote Alaskan soil sample (Rodríguez et al., 2017). In addition, we found a clade of 18 Antarctic BLAs related to CRD3-1 (77%–84.19% identity), a BLA previously found in *Erythrobacter* sp. that displays high levels of imipenem-hydrolyzing activity (Gudeta et al., 2016). Also, we observed a putative novel family constituted by a monophyletic clade sharing a common ancestor with SIE-1, first identified in *Sphingobium indicum*, which hydrolyzes carbapenems and cephalosporins and has the peculiarity of having a Glu residue in the active site (EHH/DHH active-site motif) (Wilson et al., 2021). An additional clade comprised mostly Antarctic BLAs related to FEZ-1, identified initially in environmental *Flavobacterium* species. Finally, the B2 Antarctic BLA showed a considerable distance from the closest reference (CphA-11), also corresponding to a putative novel family. All this evidence supports Antarctic and Subantarctic soils as a rich reservoir of BLAs with unknown properties.

### Relative abundance, geographical distribution, and taxonomic origin of the Antarctic and subantarctic beta-lactamases

Subsequently, the abundance of BLA genes present in the 81 metagenomes was analyzed to determine their geographical distribution. Overall, across all the sites, MBLs accounted for 53% of the total abundance, while SBLs made up 47%. The most abundant BLAs corresponded to subclass B3 (39.42%), followed by class A (28.09%), class D (14.40%), and subclass B1 (13.41%) (Figure 5A). AP and SI soils exhibited a higher total BLA abundance compared to CD (Figure 5B). At the class level, A, C, and B1 maintain this trend (Figures 5C, 5D, and 5G). Meanwhile, in Class D, B2, and B3, no significant differences were observed among the three sites (Figure 5E, 5F, and 5H).

**Figure 5.**
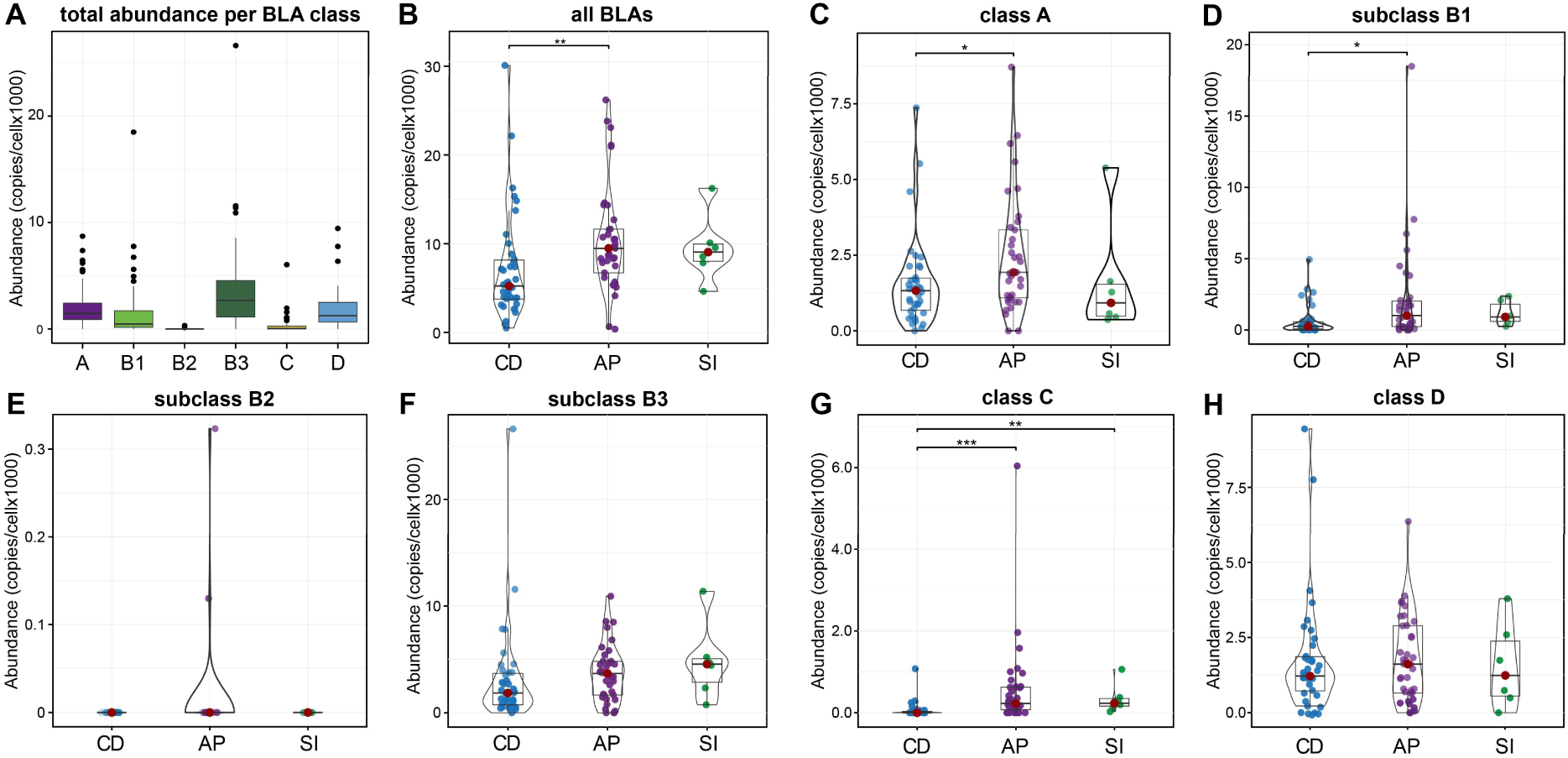
Relative abundance of Antarctic beta-lactamases across different sites. Distribution of BLA abundances (copies/cell) across cold desert (CD), Antarctic Peninsula and surrounding islands (AP), and subantarctic islands (SI) soils. A. Total BLA abundance per class across all Antarctic sites, statistical differences were calculated using the non-parametric Friedman test. ***pFDR-adjusted<0.001 (Durbin-Conover’s pairwise test). B. Total BLA abundance per Antarctic site, statistical differences were calculated using the non-parametric Kruskal-Wallis test **pFDR-adjusted<0.01 (Dunn’s pairwise test) . C-H. Total BLA abundance by class per Antarctic site, statistical differences were calculated using the non-parametric Kruskal-Wallis test *pFDR-adjusted<0.05, **pFDR-adjusted<0.01, ***pFDR-adjusted<0.001 (Dunn’s pairwise test).

We investigated the taxa associated with the Antarctic BLAs using two approaches. In a first approach, the metagenomic reads mapping to the BLAs were extracted and taxonomically classified with Single M (Figure 6A). Regarding SBLs, Class A beta-lactamases were primarily associated with the phyla Acidobacteriota (class Blastocatellia), Verrucomicrobiota (class Verrucomicrobiae), and Bacteroidota (class Bacteroidia). Class C sequences were mostly linked with Pseudomonadota (mainly Gammaproteobacteria), and Verrucomicrobiota (class Verrucomicrobiae). Meanwhile, Class D Antarctic beta-lactamases were mainly associated with Bacteroidota (class Bacteroidia), Pseudomonadota (class Alphaproteobacteria), and Acidobacteriota (Blastocatellia) (Figure 6A).

**Figure 6.**
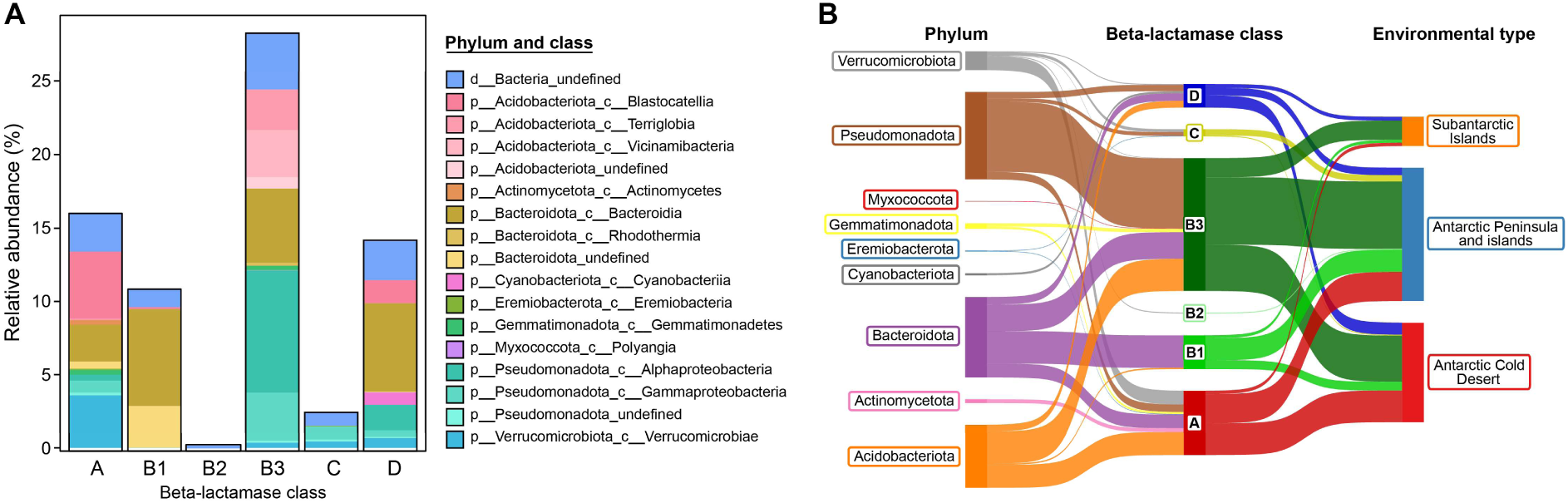
Taxonomic distribution of Antarctic beta-lactamases. A: taxonomic assignment of beta-lactamase-coding genes based on the metagenomic reads. The total relative abundance across all the sites is shown. B: Sankey plot of beta-lactamases and their host taxa detected in assembled contigs, grouped by environmental types.

For MBLs, subclass B1 was primarily associated with phylum Bacterioidota (class Bacteroidia), subclass B2 with undetermined phyla, and subclass B3 with phyla including Pseudomonadota (Alpha and Gammaproteobacteria), Acidobacteriota (classes Terriglobia and Vicinamibacteria), and Bacteroidota (Bacteroidia) (Figure 6A).

The relative abundance of the different SBL and MBL Antarctic beta-lactamases, along with their predicted source taxa, is depicted in the trees shown in Figures 3 and 4, respectively. For class A, the highest abundance was observed for proteins from Acidobacteriota sharing a common ancestor with the ASU1-1 beta-lactamase identified from functional metagenomic libraries constructed with DNA from environmental samples (Ren et al., 2023). Also, the possible novel families of Bacteroidota sharing ancestry with CME-3 and CME-4, and those with RATA-1 and VEB-9. Additionally, there is a remarkably high abundance of beta-lactamases from possible novel families hosted by Verrucomicrobiota and, to a lesser extent, Pseudomonadota.

In the case of Class C, the highest abundance was observed for Antarctic BLAs mainly from Pseudomonadota sharing ancestry with UCB-1, identified from a functional metagenomics library from agricultural and plant-associated soil (Torres-Cortés et al., 2011), and LYL-1 from *Lysobacter lactamgenus*, a bacterium known for cephalosporin biosynthesis (Kimura et al., 1996). For class D, the highest abundance was observed for OXA-like proteins from Pseudomonadota and NOD-1 from Cyanobacteriota (*Nostoc* sp.). Other Class D beta-lactamases found in high abundance corresponded to enzymes related to CPD-1 from *Chitinophaga* sp. (Bacteroidota) and OXA-347 and OXA-209 found in *Bacteroides* spp., according to the BLDB metadata.

For MBLs, it was noted that there is a class within subclass B3 corresponding to sequences from the *Acidobacteriota* phylum. We observed a high abundance of B1 enzymes, mainly from Bacteroidota, sharing ancestry with CAM-1, a carbapenemase found in *P. aeruginosa* but proposed to originate in Bacteroidota (Berglund et al., 2021; Boyd et al., 2019). Also, a high abundance of enzymes sharing ancestry with CHM-1, CEMC19-1, and MYO-1, from Bacteroidota. In contrast, the B2 beta-lactamase, distantly related to CphA, was found in very low abundance. For B3, the highest abundances were observed for beta-lactamases from Pseudomonadota, Verrucomicrobiota, and Acidobacteriota, sharing ancestry with FEZ-1, B3SU2-1, and AM1-1. Also, several enzymes related to LRA12-1 (most from Bacteroidota). Also, a high abundance of enzymes from Acidobacteriota and Pseudomonadota that are relatively close to SIE-1 and THIN-B, and others related to ALG11 from Gemmatimonadota. Also, we found some from Pseudomonadota that are distantly related to CRD3-1.

### Antarctic beta-lactamases diversity and origin according to assembled contigs and reconstructed genomes

To further investigate the Antarctic BLA genes and their genetic context, we analyzed the BLAs encoded in the contigs and MAGs recovered from the 81 metagenomes. A total of 36,811,948 contigs were assembled with a N50 median of 1143 bp (Inter Quartile Rate: 946-1385 bp) and a median largest contig of 127,219 bp (IQR: 79,610-262,104 bp). The binning of these contigs resulted in 3,576 bins with an N50 median of 5399.5 bp (IQR: 3941-9446 bp) and a median largest contig of 24,047 bp (IQR: 12,591-44,549 bp). MAG’s taxonomy coincided with almost all the previous taxa found in the unassembled reads with SingleM. Some of these MAGs correspond to poorly described taxa, including LLD-282 and JABXHR01 (Pseudomonadota, Thiohalobacterales), or Nitriliruptorales, Acidothermales, and ATN3 (Actinomycetota), among others (Table S4).

A total of 470 contigs and 175 MAGs containing beta-lactamases were identified (Figure S7). Following the relative abundance trend for each beta-lactamase class in the reads, both contigs and MAGs show a higher count of Class B3 and Class A BLAs. Regarding the taxonomic distribution (Figure 6B), it was observed that within the SBLs, Class A is associated with the highest number of different phyla, mainly Acidobacteriota, Bacteroidota, and Verrucomicrobiota. Class C is primarily associated with Pseudomonadota, Verrucomicrobiota, and Eremiobacteriota, while Class D is associated with Bacteroidota, Acidobacteriota, and Pseudomonadota. Some taxon-specific associations were observed, such as Class A and Class D being the only ones present in Actinomycetota and Cyanobacteriota, respectively.

For MBLs, subclass B3 is associated with the highest number of distinct phyla, with the main ones being Pseudomonadota, Acidobacteriota, and Bacteroidota. Subclass B1 is only associated with Bacteroidota and Acidobacteriota, while subclass B2 is exclusively associated with Verrucomicrobiota. It was noted that the phylum Verrucomicrobiota is linked to the highest number of beta-lactamases from different classes, encompassing 5/6 subclasses, with only subclass B1 missing. Similarly, Pseudomonadota, Acidobacteriota, and Bacteroidota were associated with 4/6 classes and subclasses. Regarding their geographical distribution (Figure 6B), it was observed that the contigs containing class A, B1, B3, and D beta-lactamases originate from samples across all three regions. However, those containing class C are only found in samples from AP and CD, and subclass B2 is exclusively associated with samples from AP. Notably, this region had the six known classes and subclasses of BLAs.

In the MAGs containing beta-lactamases (Figure 7), it was observed that class A is distributed across the highest number of different phyla, with the main ones being Verrucomicrobiota, Acidobacteriota, and Bacteroidota. Class C was only associated with Pseudomonadota and Verrucomicrobiota, while class D was associated with five different phyla, with the main ones being Acidobacteriota, Pseudomonadota, and Bacteroidota. Within the MBLs, subclass B3 again showed the most diverse phylum distribution, with Pseudomonadota, Acidobacteriota, and Bacteroidota being the predominant phyla. Subclasses B1 and B2 were associated only with Bacteroidota and Acidobacteriota, respectively. Interestingly, some phyla exclusive to each class were identified, which were not previously identified in the contigs. A Planctomycetota MAG was identified for class A, and a Myxococcota MAG for class B3.

**Figure 7.**
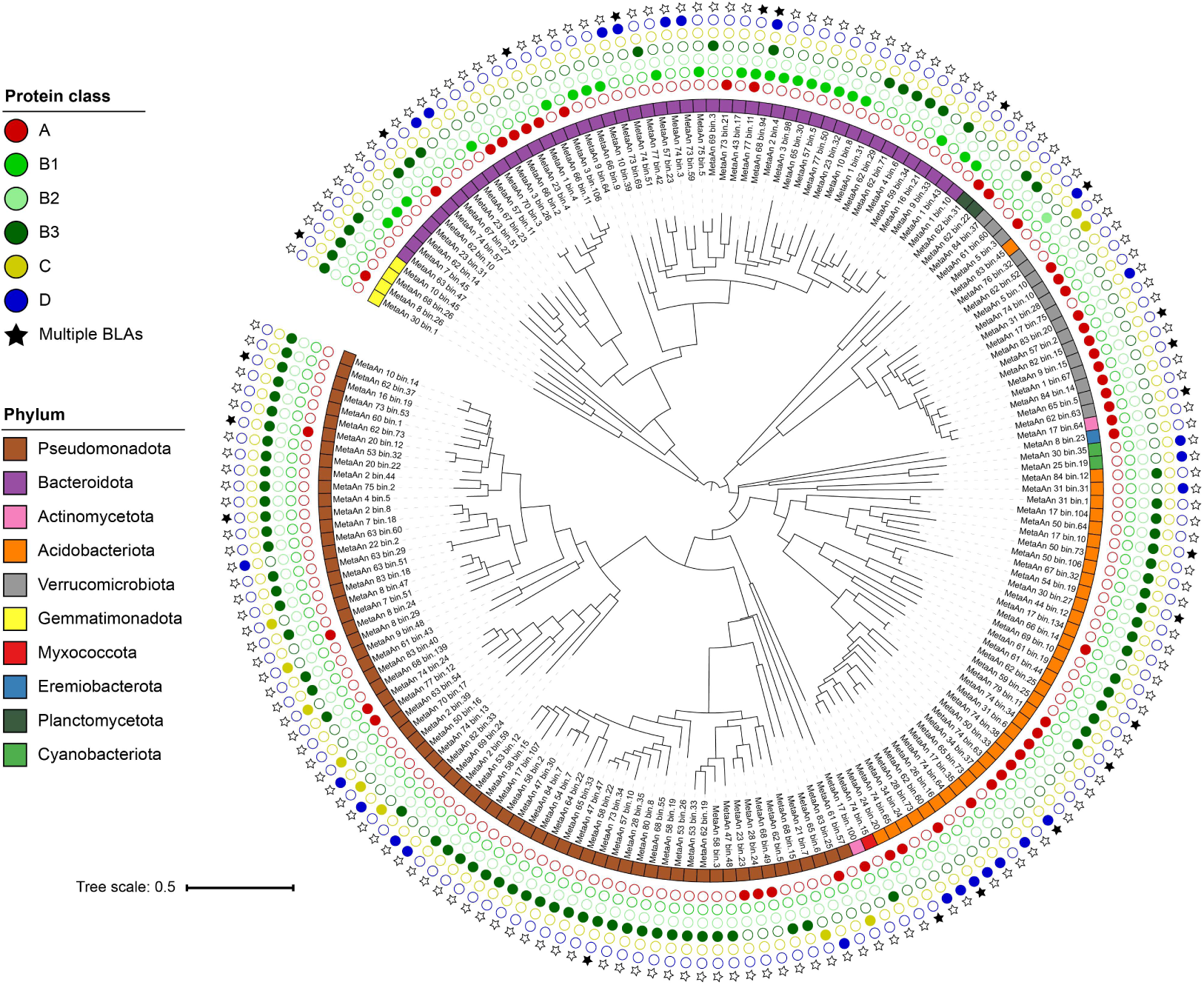
Phylogenetic relationship between assembled MAGs containing beta-lactamases. Midpoint-rooted phylogenetic tree inferred from the multiple sequence alignment of the Genome Taxonomy Database (GTDB) bacterial markers. The phylum and BLAs detected for each MAG are shown. Multiple BLA hits were highlighted with a black star.

**Figure 8.**
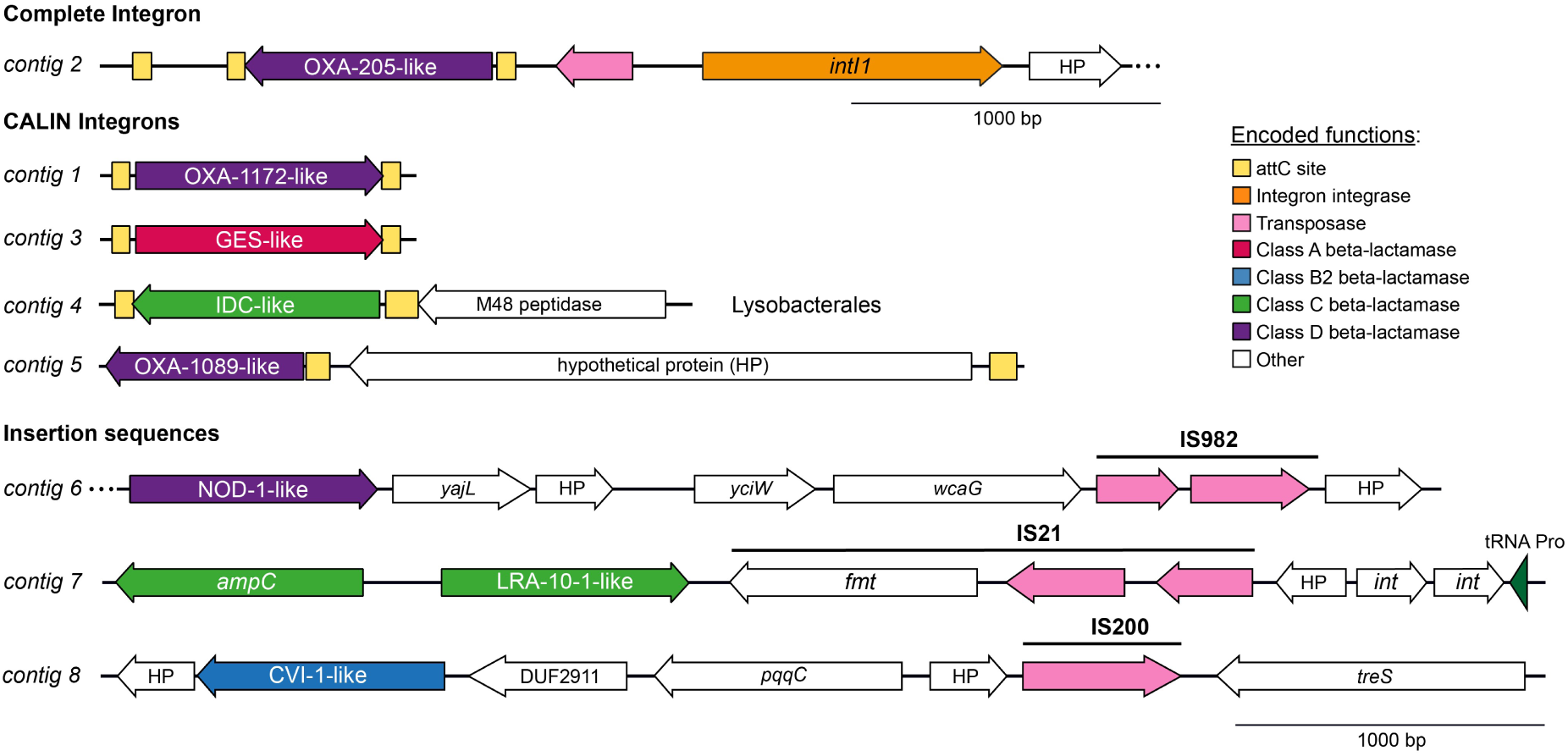
Antarctic beta-lactamases associated with mobile genetic elements. Schematic representations of mobile genetic elements associated with BLAs observed in assembled contigs of Antarctic metagenomes. Class A BLAs are shown in red, class B2 in light blue, class C in light green, class D in purple, integron integrase genes in orange, transposase genes in pink, and other unrelated genes in white. attC sites are shown as yellow boxes and tRNA genes as dark green triangles.

Resembling the findings observed with the contigs, MAGs associated with Pseudomonadota, Bacteroidota, Acidobacteriota, and Verrucomicrobiota were identified as the phyla harboring the highest number of different beta-lactamase classes (four out of six classes each). More specifically, certain MAGs were found to encode multiple BLA proteins, either from the same or different classes. Notably, the MAGs MetaAn74.57 (p Bacteroidota; g Chryseolinea) and MetaAn8.26 (p Gemmatimonadota; f Gemmatimonadaceae; g Fen-1247) encode two BLA sequences, each belonging to distinct classes (A and B3). Finally, it is worth noting that MAGs affiliated with the same genetic clades tend to harbor beta-lactamases from the same structural classes, suggesting a taxon-specific distribution pattern of BLA classes among these environmental bacteria.

### Antarctic beta-lactamases associated with mobile genetic elements

We aimed to evaluate possible BLA genes associated with mobile genetic elements, and thus, more prone to disseminate through horizontal gene transfer. We found one SBL gene encoding a class D OXA-205-like enzyme in a complete integron observed in a Pseudomonadota contig from the Burkholderiaceae family, and four BLA genes in CALIN-type integrons, one corresponding to a class D OXA-1172 type. The remaining genes likely correspond to new variants of SBL beta-lactamases: one class A (62.5% identity against GES-41) observed in a Pseudomonadota contig from the *Lysobacter* genus, one Class C (83.5% identity against IDC-1) also in a *Lysobacter* contig, and one Class D (52% identity against OXA-1089) in an Acidobacteriota contig from the Pyrinomonadales order.

In the analysis of beta-lactamase genes located near insertion sequence sequences, one Class D BLA gene (47.4% identity against NOD-1) was observed in the vicinity of an IS982 in an Acidobacteriota contig from the Pyrinomonadales order. Also, a class C BLA gene (61.2% identity against LRA10-1) was observed in the vicinity of an IS21 in a Pseudomonadota contig from the Alphaproteobacteria class. Additionally, a subclass B2 BLA gene (61.9% identity against CVI-1) was observed in the vicinity of an IS200/IS605 in a Verrucomicrobiota contig from the Chthoniobacterales order. These results suggest the potential for genetic mobilization of both well-known beta-lactamases, such as OXA-205 and OXA-1172, as well as new types of beta-lactamases, primarily of the SBL-type.

No association between predicted prophages and BLAs was observed, suggesting that potential mobilization of BLAs through means of transduction is less frequent. However, the limited length of the assembled metagenomic contigs likely leads to an underestimation of the MGEs present in the sampled soils. In a similar direction, putative plasmid sequences were not evaluated, due to most bioinformatics pipelines used to predict plasmid sequences in metagenomic samples have high false-positive rates (Pradier et al., 2021).

Together, these results suggest that at least part of the Antarctic and Subantarctic beta-lactamases are associated with mobile genetic elements and thus are more prone to be disseminated through horizontal gene transfer. More studies are required to evaluate the functional properties of these enzymes and their ability to confer beta-lactam resistance if acquired by bacterial pathogens.

## DISCUSSION

The emergence of novel beta-lactamases and their rapid dissemination via plasmids and other MGEs represents a critical escalation in the global antimicrobial resistance (AMR) crisis (Sun, 2025; Yahav et al., 2020). These enzymes, increasingly diverse and potent, can hydrolyze a broad spectrum of β-lactam antibiotics—including carbapenems—rendering many frontline treatments ineffective. Here, we showed that Antarctic and subantarctic soil microbiota stand as prominent sources of BLAs from different classes, particularly MBLs.

When investigating beta-lactamases from environmental bacteria that are phylogenetically distant from those in clinical databases, relying solely on primary-sequence-based bioinformatic identification poses the risk of false positives, particularly for sequences with low identity that fall into the so-called “twilight zone” (Rost, 1997). This is especially relevant given that many distant beta-lactamase orthologs may actually correspond to proteins with alternative functions within their respective superfamilies, such as PBP-like and MBL-fold proteins (Fröhlich et al., 2021). Since tertiary structure is generally more conserved than primary sequence, several approaches based on three-dimensional similarity have been developed to improve the identification of remote orthologs (Köstlbacher et al., 2024; Ruperti et al., 2023). In parallel, methodologies based on functional similarity have also been shown to recover orthologs that are highly divergent at the sequence level (Boger et al., 2025; Koehler Leman et al., 2023). Building upon these principles, we developed the PROSAFF pipeline, which integrates sequence, structural, and functional similarity for beta-lactamase identification. This approach enabled the recognition of phylogenetically distant Antarctic beta-lactamases that largely retained the key catalytic residues required for function, while clearly distinguishing them from members of the broader PBP-like and MBL-fold superfamilies.

Analysis of the structural class distribution revealed that class B3 and class A BLAs exhibited the highest relative abundance (39.42% and 28.09%, respectively), as well as the highest number of unique sequences (n=665 and 512, respectively). This observation contrasts with a previous study based on metagenomes from microbial mats collected on King George Island, where class C beta-lactamases were the most abundant (46.8%), followed by class B (35.5%) (Azziz et al., 2017). It has been reported that microbial mats possess a different and less diverse taxonomic composition than soil environments, which may explain these observed differences (Finke et al., 2019; Lavoie et al., 2017). Another study conducted on Russian cryosol soil samples also reported a predominance of class-C (52–83%), followed by class B BLAs (average of 14%) (Rigou et al., 2022). However, in a broader study analyzing 232 metagenomes from ten distinct environments—including glaciers, oceans, freshwater, non-agricultural soils, agricultural soils, and aerobic/anaerobic wastewater reactors—class B beta-lactamases were the most abundant overall, except for bovine and human fecal samples, where class A predominated (Gatica et al., 2019). These findings are consistent with those reported in a previous work, in which the authors analyzed 10,000 metagenomic samples and reported a high abundance of class B3 beta-lactamases in non-human-associated environments such as freshwater, lentic water, marine water, soil, and rhizosphere (Inda-Díaz et al., 2023). In contrast, class A beta-lactamases exhibited a more homogeneous distribution, with relatively higher abundance in human-associated environments.

Nonetheless, the abundance of class B beta-lactamases in human-derived samples may be on the rise, as suggested by recent studies (Jia et al., 2024), in response to the increased clinical use of beta-lactam antibiotics that are not efficiently hydrolyzed by class A enzymes. This is particularly relevant given reports of class B beta-lactamases originating from environmental bacteria—particularly those of subclass B3, as identified in the present study—that are capable of hydrolyzing all beta-lactam antibiotics (Krco et al., 2023). These findings highlight Antarctic environmental bacteria as a potential significant reservoir of previously uncharacterized beta-lactamase variants, some of which may exhibit increased hydrolytic activity against clinically relevant beta-lactam antibiotics.

In terms of taxonomic distribution, consistent with previous studies, beta-lactamases were predominantly associated with the phyla Pseudomonadota and Bacteroidota, regardless of enzyme class or sample origin (Berglund et al., 2017; Inda-Díaz et al., 2023). Notably, we also unveiled an unprecedented diversity of BLAs from members of the phylum *Acidobacteriota*, being the first large-scale BLA screening to report such an association, particularly relevant for class A beta-lactamases. Although *Acidobacteriota* primarily comprises uncultivable representatives, a subclass B2 beta-lactamase (CAM-2) has previously been reported within the chromosome of a metagenome-assembled genome (MAG) affiliated with the family *Pyronomonadaceae* of the phylum *Acidobacteriota*, recovered from sediments in Shenzhen Bay, China (Xiang and Li, 2022). This enzyme was proposed as a potential ancestor of the clinically disseminated CAM-1 variant found in *Pseudomonas aeruginosa*. In the present study, 57 sequences were identified with 52–83% sequence identity to CAM-1, suggesting that Antarctic environmental *Acidobacteriota* may also act as reservoirs for novel beta-lactamase variants of clinical interest.

Although most beta-lactamases identified in this study are chromosomal, five were found to be associated with either complete or partial integrons, and three were located in proximity to insertion sequences. Notably, the OXA-205 beta-lactamase variant was identified, which has previously been reported in a class 1 integron from a clinical *Pseudomonas aeruginosa* isolate (Krasauskas et al., 2015), as well as in Antarctic microbial mats (Antelo et al., 2021), microplastic-associated samples along watersheds, and activated sludge from wastewater treatment plants in China (Li et al., 2022; Zhang et al., 2021). Additionally, two BLA genes previously associated with clinical or human-impacted environments were identified here for the first time in a pristine, non-human-associated setting: the class D variant OXA-1171/OXA-544, initially described in clinical *P. aeruginosa* isolates, and a class C beta-lactamase sharing 83.1% identity with IDC-1, an integron-derived cephalosporinase previously detected only in wastewater samples from Asia (Böhm et al., 2020; Laudy et al., 2017).

The remaining beta-lactamases exhibited less than 65% sequence identity to reference sequences in the Beta-Lactamase DataBase (BDLB), indicating that they may represent previously uncharacterized families. This highlights the need for further functional characterization and assessment of their potential for horizontal gene transfer, particularly given that many beta-lactamases acquired by clinical pathogens have originated from environmental Proteobacteria (Berglund et al., 2017; Ebmeyer et al., 2021; Gholipour et al., 2024), the same phylum to which the potentially mobile beta-lactamases identified in this study belong.

Studies of the Antarctic resistome have revealed that this environment harbors a remarkable diversity of bacteria and natural antibiotic resistance genes across different families, despite limited human impact (Arros et al., 2024; Marcoleta et al., 2022; Van Goethem et al., 2018). It has been proposed that such resistance genes likely evolved in response to interactions with Antarctic microorganisms capable of producing antibiotics (Van Goethem et al., 2018). In particular, beta-lactamases from Antarctic bacteria may represent a defense mechanism against beta-lactam–producing fungi commonly found in certain soil samples (Pantůček et al., 2018). Alternatively, they might fulfill physiological functions unrelated to antibiotic resistance—for instance, SBLs could contribute to cell-wall biosynthesis (Jacobs et al., 1994; Massova and Mobashery, 1998), while MBLs may be involved in quorum-sensing (Fisher and Mobashery, 2023; Selleck et al., 2020). The latter hypothesis is especially compelling, given that Antarctic soils have been shown to contain a broad diversity of homoserine lactone (HSL)–based quorum-sensing systems, which may mediate cell-to-cell communication as an adaptive strategy in extreme environments (Wong et al., 2019). Further studies are required to elucidate the natural functions of beta-lactamases identified in Antarctic soil bacteria.

Advances in next-generation sequencing (NGS) technologies and tools for identifying antibiotic resistance genes have greatly expanded our knowledge of novel gene families and associated bacterial lineages, surpassing what is known from clinical settings (Larsson and Flach, 2022). This has been particularly significant for beta-lactamase genes, for which it has been estimated that over ∼100,000 remain to be discovered. Consequently, there is a growing emphasis on functional and structural characterization of bioinformatically predicted genes from natural environments (Bush, 2023; Gholipour et al., 2024). In this context, the discovery of beta-lactamase genes in a low-impact environment such as Antarctica provides an opportunity to explore the evolution of enzymatic activity under reduced selective pressure compared to clinical settings—likely shaped instead by natural production of beta-lactams by co-occurring microorganisms (Gholipour et al., 2025)—or potentially linked to alternative functions within their superfamilies (Fröhlich et al., 2021). Furthermore, several beta-lactamases identified here may represent models of cold-adapted proteins, as they originate from bacteria inhabiting environments with winter temperatures as low as –60 °C (Turner et al., 2020). Understanding the structural and kinetic bases of cold adaptation is highly relevant to industrial applications, since cold-active enzymes can reduce energy costs by operating without heating, while lowering the carbon footprint compared with processes that require high temperatures (Kumar et al., 2021).

Overall, this study introduces a new bioinformatic methodology that enabled the identification of novel beta-lactamase variants spanning all structural classes, with a particularly high prevalence of MBLs. The structural and functional characteristics of these enzymes, as well as their potential for genetic mobilization, remain to be fully investigated. Importantly, their origin from Antarctic bacteria highlights their relevance both as models for natural antibiotic resistance mechanisms and as systems for studying enzymatic adaptation to cold environments.

## METHODS

### Antarctic soil metagenome database construction

This study included 66 metagenomes downloaded from the NCBI Sequence Read Archive (SRA) database available until September 26, 2023, plus 12 metagenomes sequenced by our research group. As inclusion criteria, we used all the metagenomes that matched the query “antarctic soil metagenomes” which had their geographical localization, effectively came from soil samples, and corresponded to shotgun metagenomic sequencing. In cases of replicates from the same soil samples, we selected the one with the most base pairs sequenced. The metagenomes FASTQ files were downloaded using their SRA run accession and prefetch and fasterq-dump tools combination from SRA-toolkit v2.11.3.

### Metagenomic reads quality assessment and trimming

Raw Illumina reads were quality-checked using fastQC v0.11.9 (Andrews et al., 2012) and processed using fastp v0.23.4 (Chen et al., 2018) with default parameters to filter low-quality reads and possible adapters. We also assessed human contamination through mapping all the metagenomes to the human reference genome GRCh38.p14 (accession: GCF_000001405.40) using bwa-mem2 v2.2.1 (Vasimuddin et al., 2019) with default parameters and samtools v1.13 (Danecek et al., 2021) to discard all the mapped reads.

### Sequence and microbial diversity analysis

Sequence reads for each metagenome were processed using SingleM pipe v0.18.3 (Woodcroft et al., 2025) with default parameters, and then summarized using the summarise subcommand to determine operational taxonomic units (OTUs) at the species level by clustering similar sequences at a 96.66% identity threshold for 43 single-copy universal bacterial and archaeal genes (SCGs). Next, for each SCG, α-diversity metrics (Richness, Pielou evenness, and Shannon Index) were determined in each metagenomic sample using custom Perl scripts. Additionally, SingleM results were used to determine the taxonomic profiles of the observed communities in each soil sample based on the GTDB r220 taxonomy database (Chaumeil et al., 2022). These profiles were used to estimate β-diversity (Bray-Curtis dissimilarity) between all the metagenomes and assess possible associations with the environmental type using the vegan (Oksanen et al., 2025) and pairwiseAdonis (https://github.com/pmartinezarbizu/pairwiseAdonis) packages in a custom R script. NonPareil v3.401 (Rodriguez-R et al., 2018) was used in kmer mode to estimate metagenomic coverage and sequencing effort for each metagenomic sample. MASH v2.3 (Ondov et al., 2016) was used to estimate k-mer distance between metagenomic samples with options sketch, dist, and default parameters. Figures were plotted using the ggplot2 and ggstatsplot (Patil, 2021) packages in R.

### Metagenome assembly and protein prediction

Metagenomic trimmed reads were assembled into contigs using metaSPAdes v3.15.5 (Nurk et al., 2017) with default parameters. These contigs were processed using Prodigal v2.6.3 (Hyatt et al., 2010) in meta mode to translate the nucleotide sequences of all the metagenomic samples. Additionally, protein sequences were directly assembled from trimmed reads using PLASS (Steinegger et al., 2019) in assemble mode with default parameters, adjusting the –min-length to 30 or 40 depending on the read length of the samples (100 bp or <150 bp, respectively).

### Beta-lactamase protein sequence database construction and identification of (sub)Antarctic beta-lactamases

A total of 7,566 BLA protein sequences were downloaded from the Beta-Lactamase database BLDB (Naas et al., 2017) on October 5th, 2023, making sure to preserve the protein family and enzyme class associated with each of them. Additionally, a total of 6 PBP-fold superfamily and 18 MBL-fold superfamily non-beta-lactamase protein sequences were added to use as true negatives (TN DB, Table S2).

BLDB protein amino acid lengths were determined, and standard statistical metrics were calculated for the obtained length distribution (Q0: 95, Q1: 284, Q2: 313, Q3: 377, Q4: 428, arithmetic mean: 329.336, standard deviation: 52.4, mode: 377). Based on the length distribution of BLDB proteins, PLASS assembled and Prodigal predicted metagenomic proteins were filtered by amino acid length (minimum length: 100 aa, maximum length: 500 aa). 98,161,403 PLASS proteins (PLASS DB) and 9,024,650 Prodigal (Meta DB) proteins were kept after applying this size filter. Proteins from each database (BLDB, PLASS DB and Meta DB) were redundancy filtered separately with MMSeqs2 v15.6f452 (Steinegger and Söding, 2017) in Linclust mode (Steinegger and Söding, 2018) using sequence identity ≥ 0.99 and bidirectional coverage ≥ 0.99 thresholds (--alignment-mode 3 --cov-mode 0 --cluster-mode 2 --realign), only keeping the representative protein sequences for further steps.

All 65,532,405 redundancy filtered protein sequences (5458 from BLDB, 58,058,386 from PLASS DB, 7,468,537 from Meta DB and all 24 from TN DB) were redundancy filtered as a single set following the same procedure described previously. Then, the resulting 62,047,905 protein set was clustered with low stringency thresholds (sequence identity ≥0.2 and bidirectional coverage ≥0.8) using MMSeqs2 as described previously. All 267 low-stringency clusters containing BLDB or TN DB proteins were kept for further analysis. 81 of these clusters also contained 3,502 Antarctic proteins (PLASS DB or Meta DB).

All 3,502 relevant Antarctic proteins selected in the previous step and 5,482 redundancy-filtered reference sequences (5,458 BLDB proteins and 24 TN DB proteins) were used as input for the atomic-level evolutionary-scale protein structure predictor ESMFold v1.0.3 (ESM-2 model) (Lin et al., 2023). Next, Foldseek v8.ef4e960 (van Kempen et al., 2024) was used to cluster all 8,984 predicted protein structure models with highly stringent global and local structural similarity thresholds (TM - Score >= 0.75 and LDDT >= 0.75) while keeping the lax sequence identity thresholds of the previous MMSeqs2 clustering (sequence identity >= 0.2 and bidirectional coverage >= 0.8) (--cov-mode 0 -e 1 --cluster-mode 2 --cluster-reassign -s 7.5 --tmalign-hit-order 3 --tmalign-fast 0 --sort-by-structure-bits 1). Then, from the 70 structural clusters obtained, 34 of them contained both reference and antarctic proteins (25 clusters contained both antarctic candidates and BLDB proteins, while 9 contained both antarctic sequences and TN DB references).

All 3,025 Antarctic protein sequences contained in the previously described 34 relevant structural similarity clusters were considered as putative beta-lactamases (2174) or putative true negative non-beta-lactamase proteins (851). Next, the enzyme class from reference proteins contained in each structural cluster was inherited to putative antarctic BLAs and TNs (no cluster contained more than one enzyme class). Similarly, redundant protein sequences from previous steps (>= 0.99 sequence identity and ≥0.99 bidirectional coverage clusters) were assigned to structural clusters by reverse inheritance. Finally, the 3,025 selected Antarctic proteins and all 7,590 reference proteins were clustered iteratively using DIAMOND BLASTP v2.1.9 (Buchfink et al., 2021) in ultra-sensitive mode until each protein belonged to a unique single-linkage cluster (99% identity and 99% bidirectional coverage thresholds). The 4,892 representatives of these clusters were selected as the final Antarctic BLA set (1,916 proteins), Antarctic true negative set (692 proteins), reference BLA set (2,260 proteins), and reference true negatives set (24 proteins).

### Multiple amino acid sequence alignment and phylogenetic analyses

For each of the 34 previously mentioned relevant structural clusters and 5 more structural clusters containing only reference True Negatives, the final protein dataset corresponding to each of those clusters was redundancy filtered with MMSeqs2 as previously described using sequence identity ≥0.9 and bidirectional coverage ≥0.9 thresholds. Then, the obtained 1,184 non-redundant protein dataset was divided into two groups: the corresponding PBP-like (677 proteins) and MBL-like (507 proteins). Next, the protein sequences in each group were aligned using MAFFT v7.505 (Katoh and Standley, 2013) in globalpair mode with 1000 iterations, followed by trimming of constant and uninformative sites according to alninfo analysis performed by IQ-TREE v2.2.5 (Minh et al., 2020). Trimmed alignments were then used as input for maximum likelihood tree inference using IQ-TREE (--seed 139808 --bnni --allnni) with 1000 ultrafast bootstraps (Hoang et al., 2018) and the ModelFinderPlus algorithm (Kalyaanamoorthy et al., 2017) to select the best aminoacid substitution model for each tree according to the BIC criterion, Q.pfam+R10 for the PBP-like superfamily tree and WAG+F+R9 for the MBL-like superfamily tree. Phylogenetic trees visualization was performed using iTOL v7.0 (Letunic and Bork, 2021).

### Beta-lactamases structural alignment and visualization

All the predicted structures for the 1,184 non-redundant proteins which were used as input for phylogenetic trees inference were structurally aligned using US-align v20240319 (Zhang et al., 2022) (-mol prot -mm 4 -ter 2). Structure PDB files were imported into PyMOL v3.1.3 (Schrodinger, LLC), then signal peptides detected by SignalP v6.0 (Teufel et al., 2022) were removed. Finally, structure visualizations cartoons were made in ray_trace_mode 3.

### Contig binning and reconstruction of metagenome-assembled genomes (MAGs)

For metagenomic contig binning, the trimmed reads were mapped to their respective contigs in all the samples using bwa-mem2. The SAM files were then filtered and sorted using samtools and processed with the jgi_summarize_bam_contig_depths command to obtain the contig coverage depth per sample. These files were required as additional input for binning the contigs using MetaBAT2 v2.15 (Kang et al., 2019). Using the default parameters, a total of 3,576 bins were obtained from these metagenomic samples. All the bins were quality checked using CheckM2 v1.0.1 (Chklovski et al., 2023) and the completeness and contamination results were used to calculate quality (Completeness -5*Contamination). Under this parameter, we identified 1250 high-quality MAGs (Quality > 50).

### Taxonomic assignment of MAGs and metagenomic contigs

Previous BLAs representatives were used to determine which contigs and MAGs contain these enzymes using DIAMOND BLASTX with default parameters. The results were filtered using a custom R script setting a minimum 70% identity and 80% coverage match, finding a total of 547 contigs and 181 MAGs containing BLAs. Taxonomic assignment of contigs containing BLAs was performed using the Contig Annotator Tool CAT v6.0 (Von Meijenfeldt et al., 2019) in sensitive mode supplemented with the GTDB r220 database (Chaumeil et al., 2022). The taxonomy and BLA class information of all the contigs were summarized in a sankey diagram plotted in R using the ggsankey package (https://github.com/davidsjoberg/ggsankey). High and low quality MAGs were also taxonomically identified using GTDB-tk v2.3.0 in classify_wf mode with GTDB r220 database. The MSA output of this workflow was used as input to IQ-TREE in autodetection mode (--seed 314716, best model: LG+F+R6, bootstrap: 1000) for maximum likelihood tree inference of the MAGs comprising BLAs. Tree visualization and editing was performed with ITOL.

### Beta-lactamase genes relative abundance calculation

The final 4,176 BLA set was divided into six groups according to the BLA class assigned to each protein (A, B1, B2, B3, C and D). Next, for each group a multiple sequence alignment (MSA) was performed using MAFFT as described previously. Each MSA was used as input to build a profile hidden Markov model (HMM) using HMMER v3.2.1 (Eddy, 2011).

Then, all BLAs of the respective class observed in taxonomically assigned contigs were aligned back into the HMM to obtain new reference MSAs. Then, each reference MSA and respective HMM was used as input for SingleM seqs v0.18.3 (Woodcroft et al., 2025) to find the most conserved 20 amino acid stretch in the HMM. Next, each HMM was used to build a GraftM package using GraftM v0.15.1 (Boyd et al., 2018). Finally, the designated position in each HMM and the corresponding GraftM package were used as input to build SingleM packages.

Next, all six built SingleM packages (one for each BLA class) were used as input to find BLA-containing sequencing reads in each metagenome using SingleM pipe mode. All identified BLA-containing reads for each metagenome were taxonomically assigned using the Read Annotator Tool RAT v6.0 in sensitive mode (Hauptfeld et al., 2024) supplemented with the GTDB r220 database and the previously taxonomically assigned contigs by CAT. Then, the specific BLA contained in each read was determined using MMSeqs2 v15.6f452 easy-search mode (-s 7 --alignment-mode 3) and the previously determined BLA database (4,176 proteins) and True Negatives database (716 proteins). Stringent parameters previously reported to search for antibiotic resistance genes (ARGs) in reads (Yin et al., 2023) were applied (e-value <= 1e-7, identity >= 80%, query coverage >= 75% and alignment length of at least 25 amino acids). False positives were discarded. Finally, the relative abundance of each BLA (copies/cell) in all metagenomes was determined according to the following previously reported equation (Yin et al., 2023):

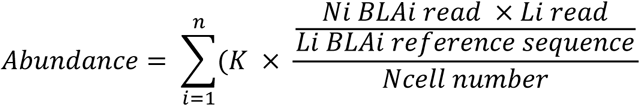

Where Ni BLAi read is the number of reads coding for a specific BLA, Li read is the read length, Li BLAi reference sequence is the length of the nucleotide sequence of the BLA, Ncell number corresponds to the estimated number of cells in the metagenome, and K is a normalization parameter for multi-gene antibiotic resistance mechanisms (for BLAs, K = 1). The number of cells in each metagenome was estimated using the SingleM microbial_fraction subcommand (Eisenhofer et al., 2024) as the quotient between the number of nucleotide bases and the average prokaryotic genome size.

### Functional prediction and similarity clustering of beta-lactamases and related proteins

For each of the 4,982 proteins in the database, functional prediction annotation based on Gene Ontology (GO) terms was performed using AnnoPRO v0.2-alpha+2.g547770e (Zheng et al., 2024). Then, functional similarity between proteins was determined by calculating the cosine distance between GO-term probability vectors, as previously done (Koehler Leman et al., 2023). Additionally, Uniform Manifold Approximation and Projection (UMAP) (McInnes et al., 2018) was performed using all 4982 GO-term vectors to determine functional similarity groups between proteins and plot results (n_components=2, random_state=42, n_neighbors=100, min_dist=0.1, metric=’cosine’).

### MGE analysis

All contigs coding for BLAs were used as input for the predictions of complete integrons, clusters of attC sites lacking integron-integrases (CALIN), and integron integrases using IntegronFinder v2.0.5 (Néron et al., 2022). Integrated prophages were predicted using the PHASTEST (Wishart et al., 2023) web server (accessed November 26th, 2024) and Phigaro v2.4.0 (Starikova et al., 2020). Complete and partial insertion sequences were predicted using ISEScan v 1.7.2.3 (Xie & Tang, 2017). Predicted ISs located <= 5000 bp upstream or downstream from BLA genes were considered relevant hits. MGEs genomic features were identified and annotated using Bakta v1.10.1 (Schwengers et al., 2021) in meta mode using the full Bakta database v5.1 (https://zenodo.org/records/10522951).

## ACKNOWLEDGEMENTS

We want to sincerely acknowledge Dr. Silvia Batista, Dr. Gastón Azziz, Dr. Matías Giménez, and Josefina Vera (Montevideo, Uruguay) for giving us access to three Antarctic soil metagenomes, and for very insightful discussions regarding Antarctic beta-lactamases.

