## Supplementary material for "A structure-guided pipeline to uncover the underexplored beta-lactamases from Antarctic and Subantarctic soil microbiota": Coche Arros et al Supplementary Material.pdf

### Supplementary Information

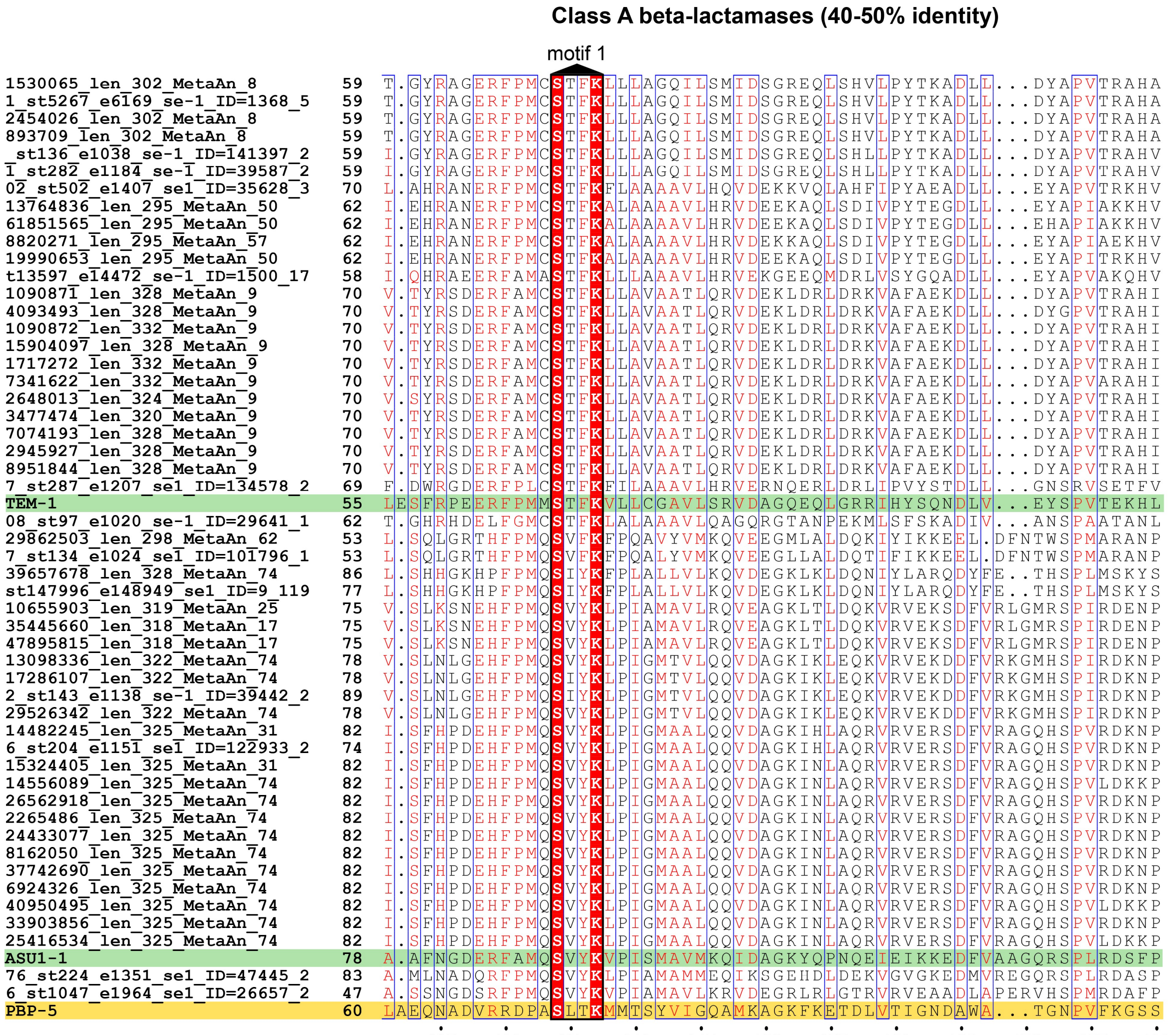


**Figure S1.** Multiple amino acid sequence alignment of key conserved regions among previously described class A beta-lactamases (shaded in green) and predicted Antarctic orthologs sharing 40-50% identity with the reference sequences. PBP-5 (shaded in yellow) was included for comparison, as it belong to the PBP-like superfamily but does not have beta-lactamase activity (true negative).

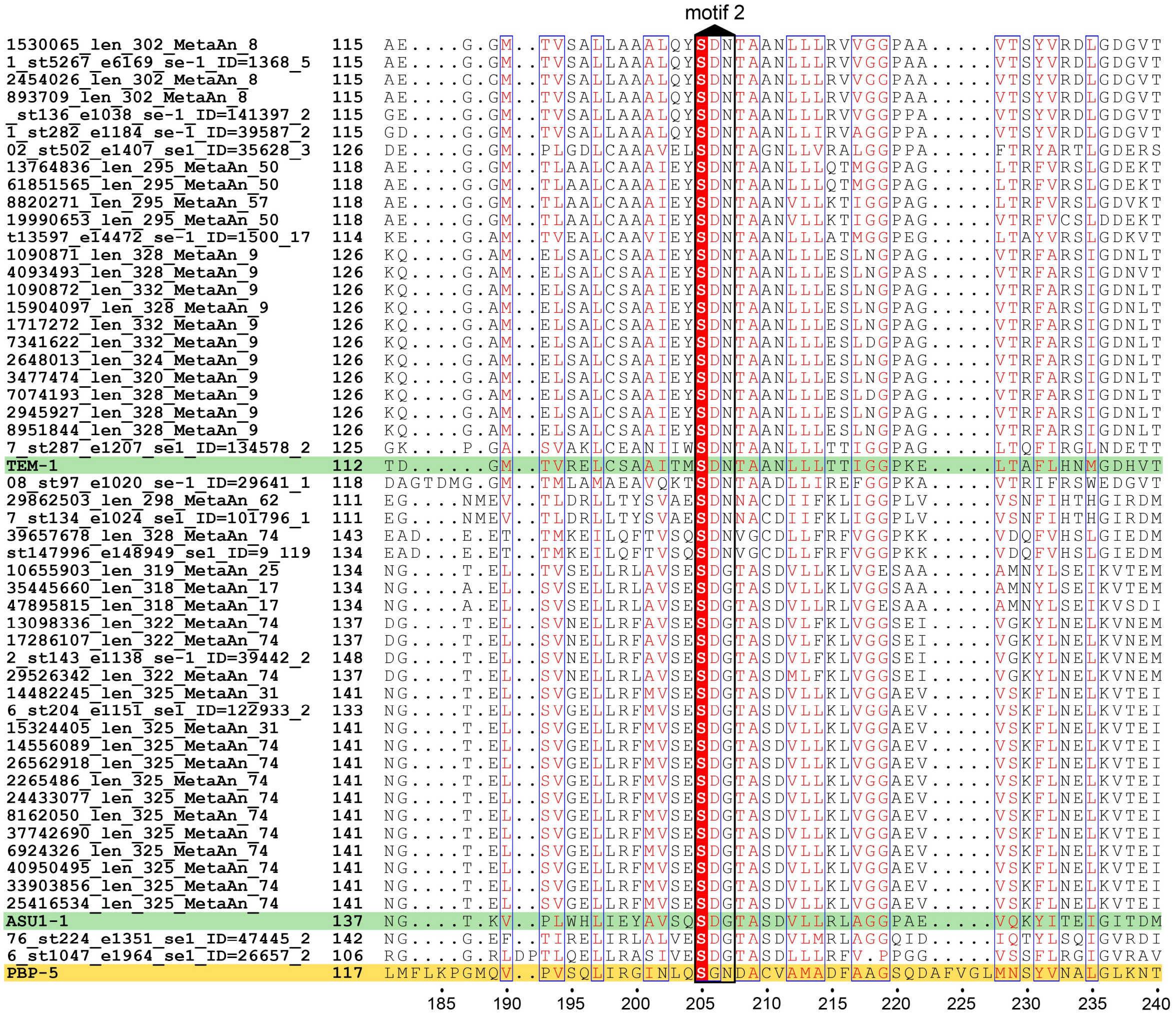


**Figure S1 (part 2/4).**

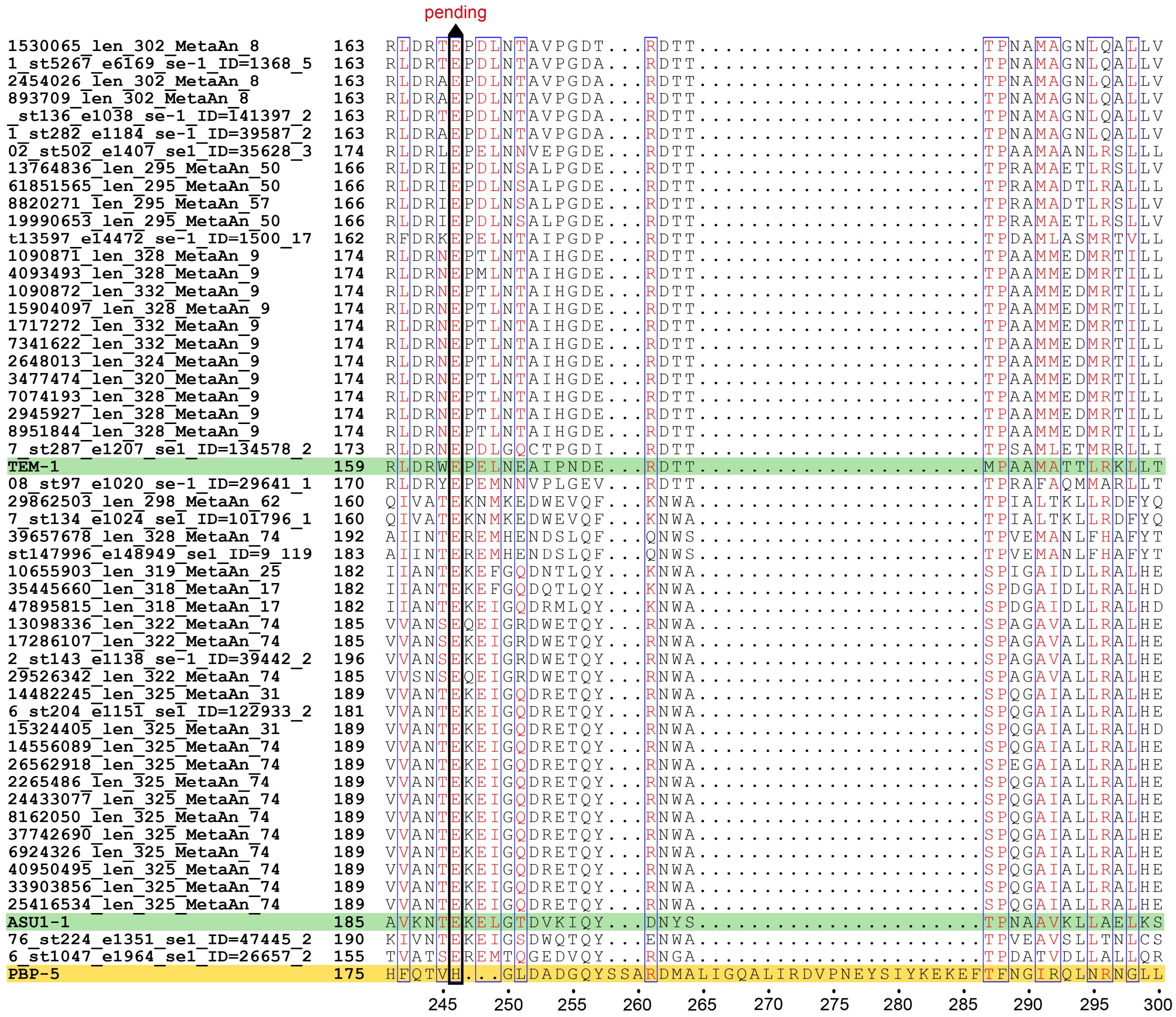


**Figure S1 (part 3/4).**

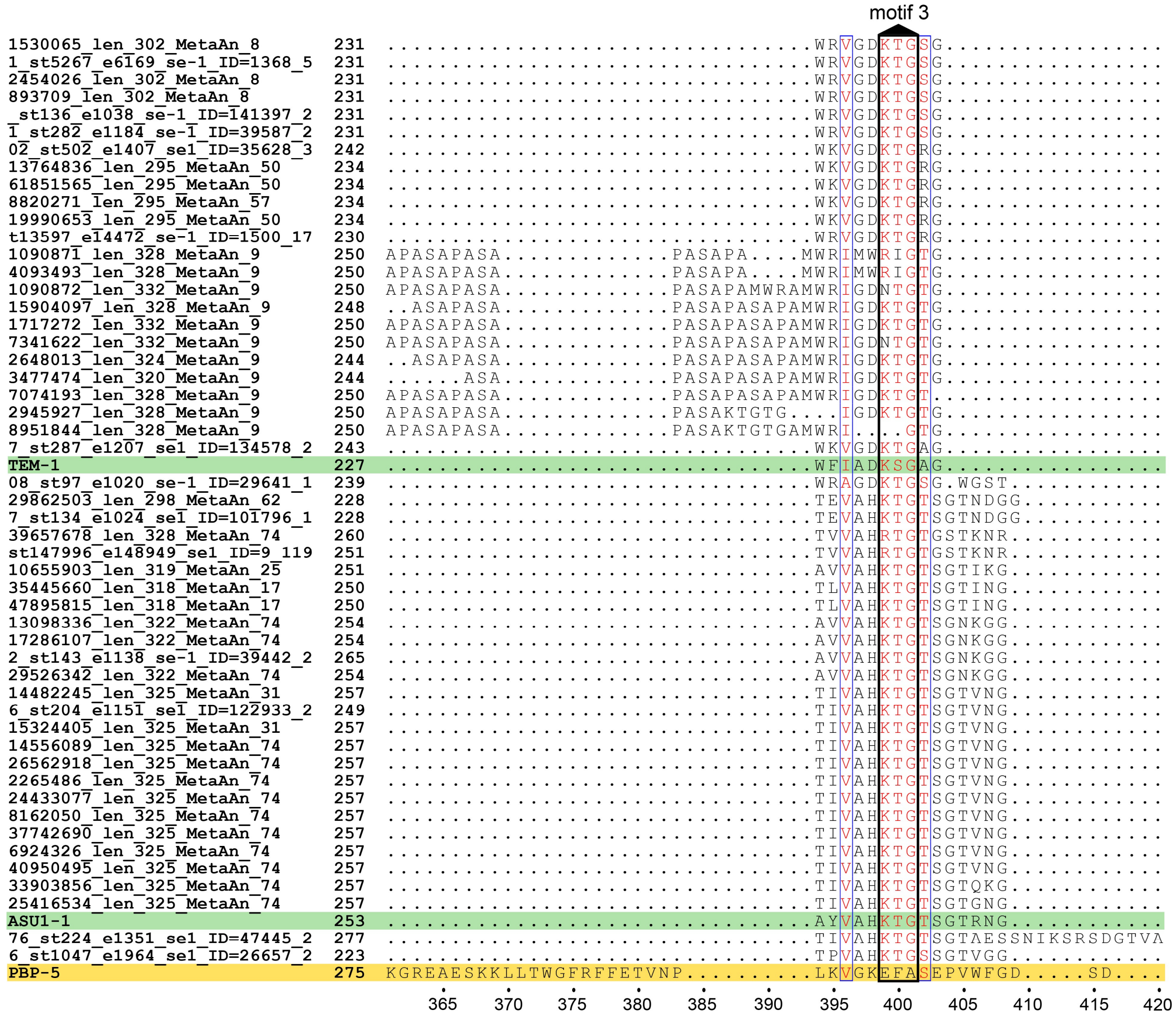


**Figure S1 (part 4/4).**

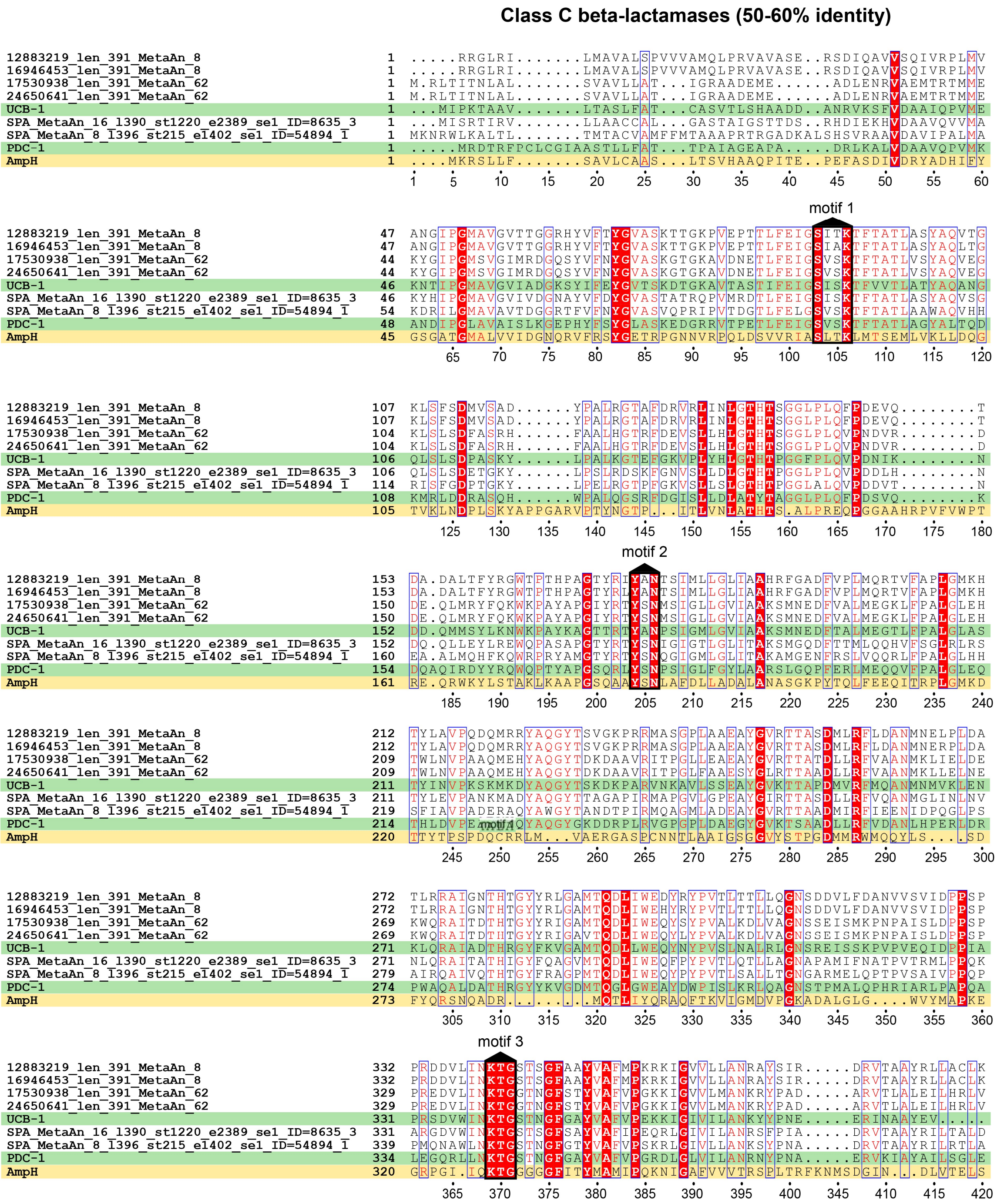


**Figure S2.** Multiple amino acid sequence alignment of key conserved regions among previously described class C beta-lactamases (shaded in green) and predicted Antarctic orthologs sharing 50-60% identity with the reference sequences. AmpH protein (shaded in yellow) was included for comparison, as it belong to the PBP-like superfamily but does not have beta-lactamase activity (true negative).

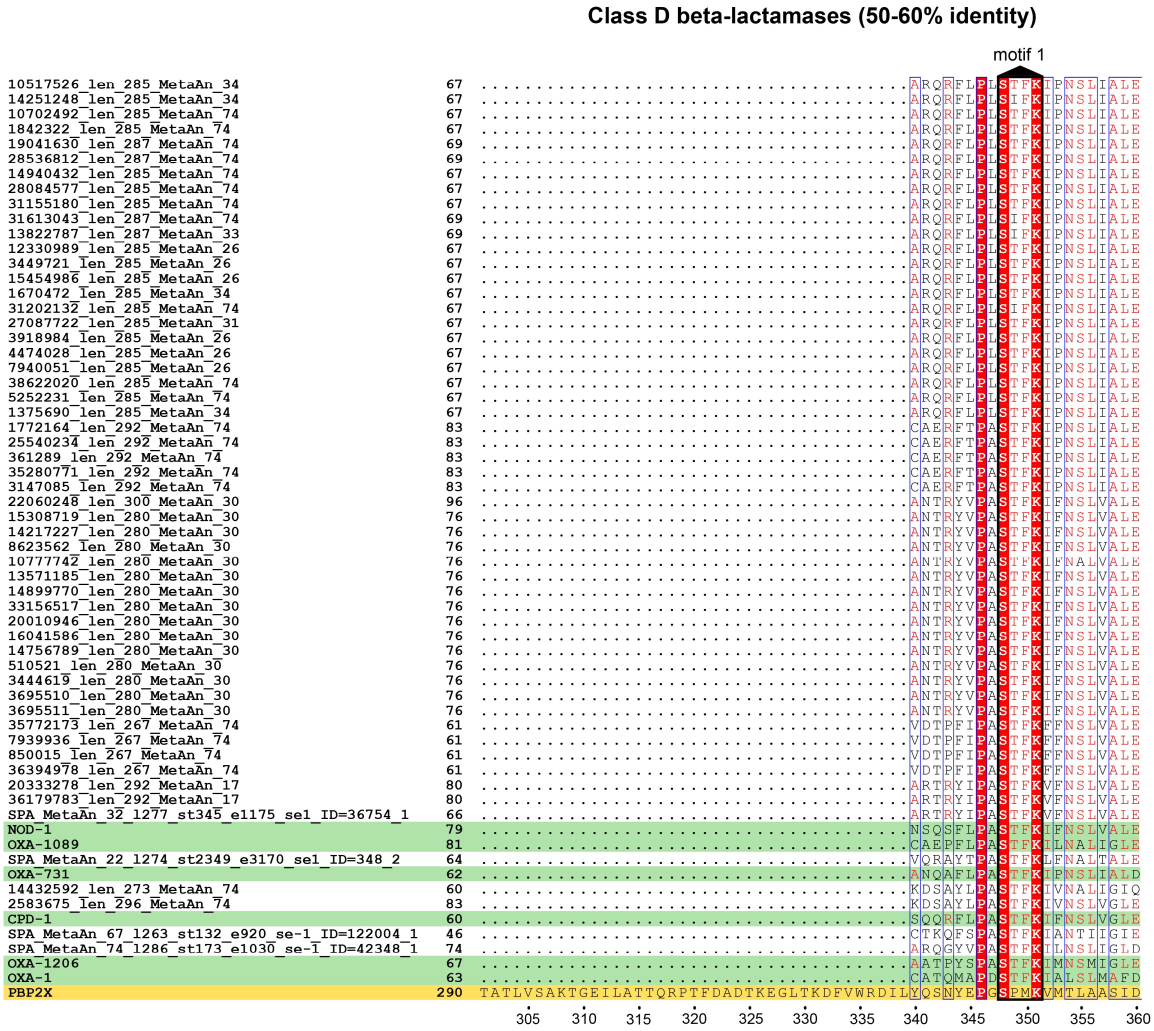


**Figure S3.** Multiple amino acid sequence alignment of key conserved regions among previously described class D beta-lactamases (shaded in green) and predicted Antarctic orthologs sharing 50-60% identity with the reference sequences. PBP2X protein (shaded in yellow) was included for comparison, as it belong to the PBP-like superfamily but does not have beta-lactamase activity (true negative).

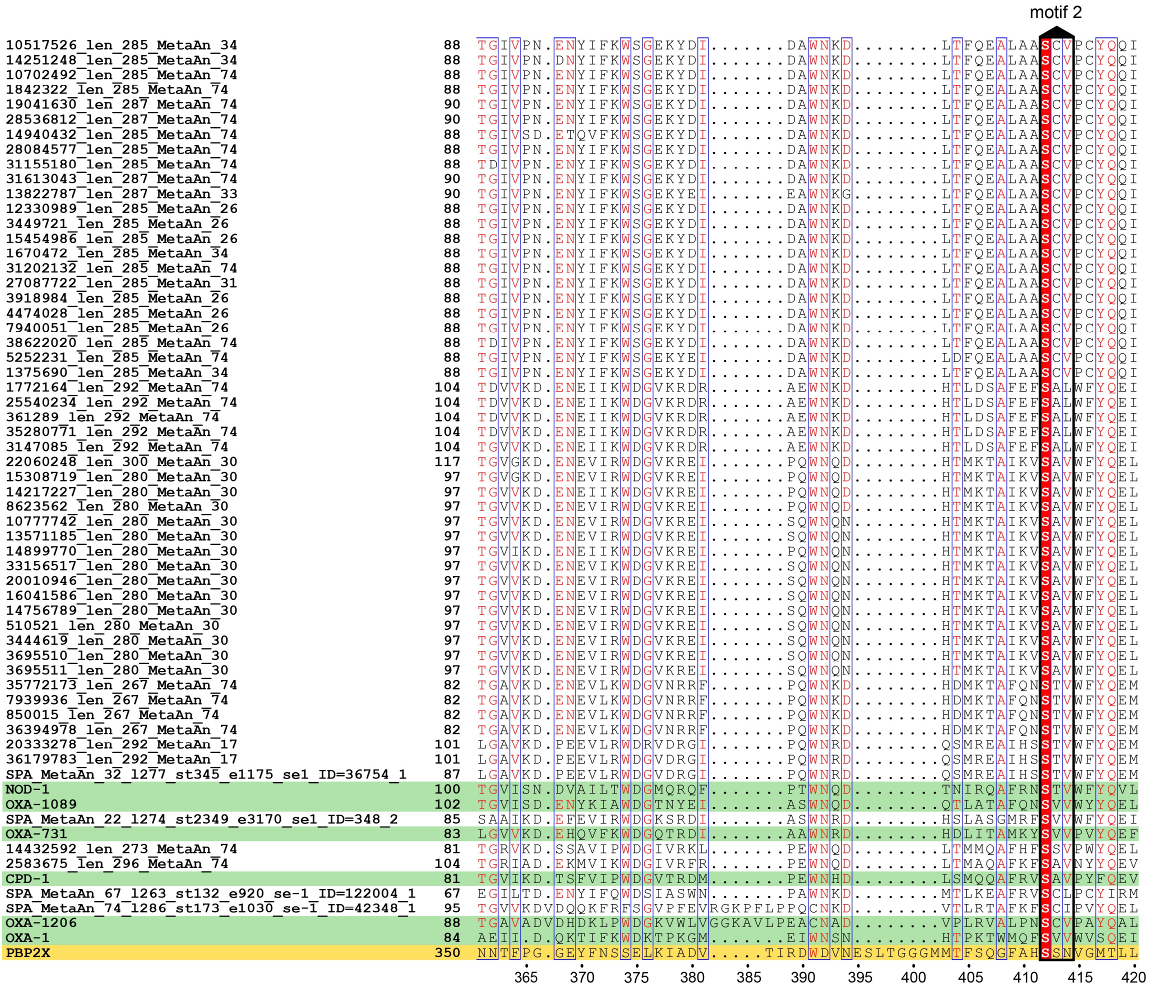


**Figure S3 (part 2/4).**

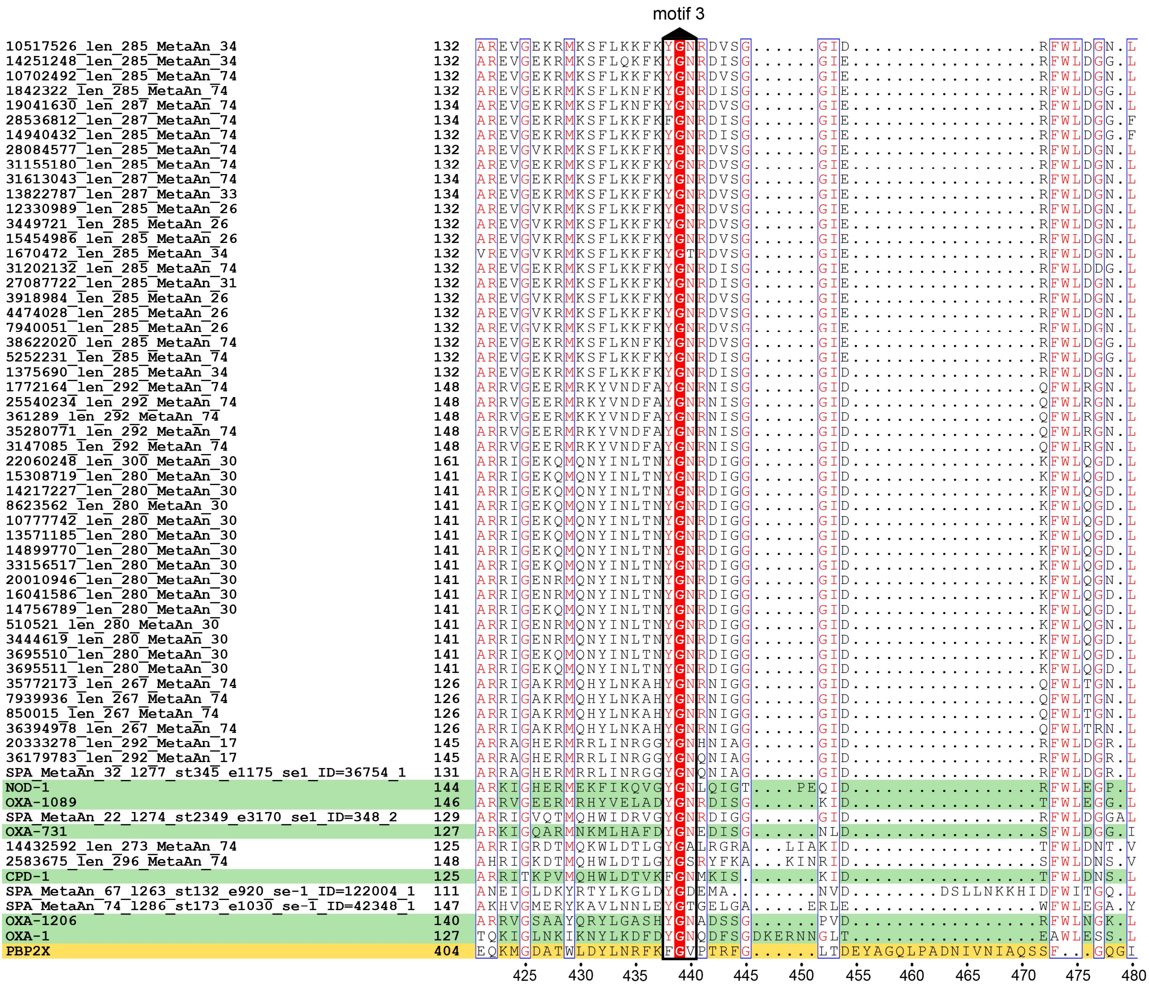


**Figure S3 (part 3/4).**

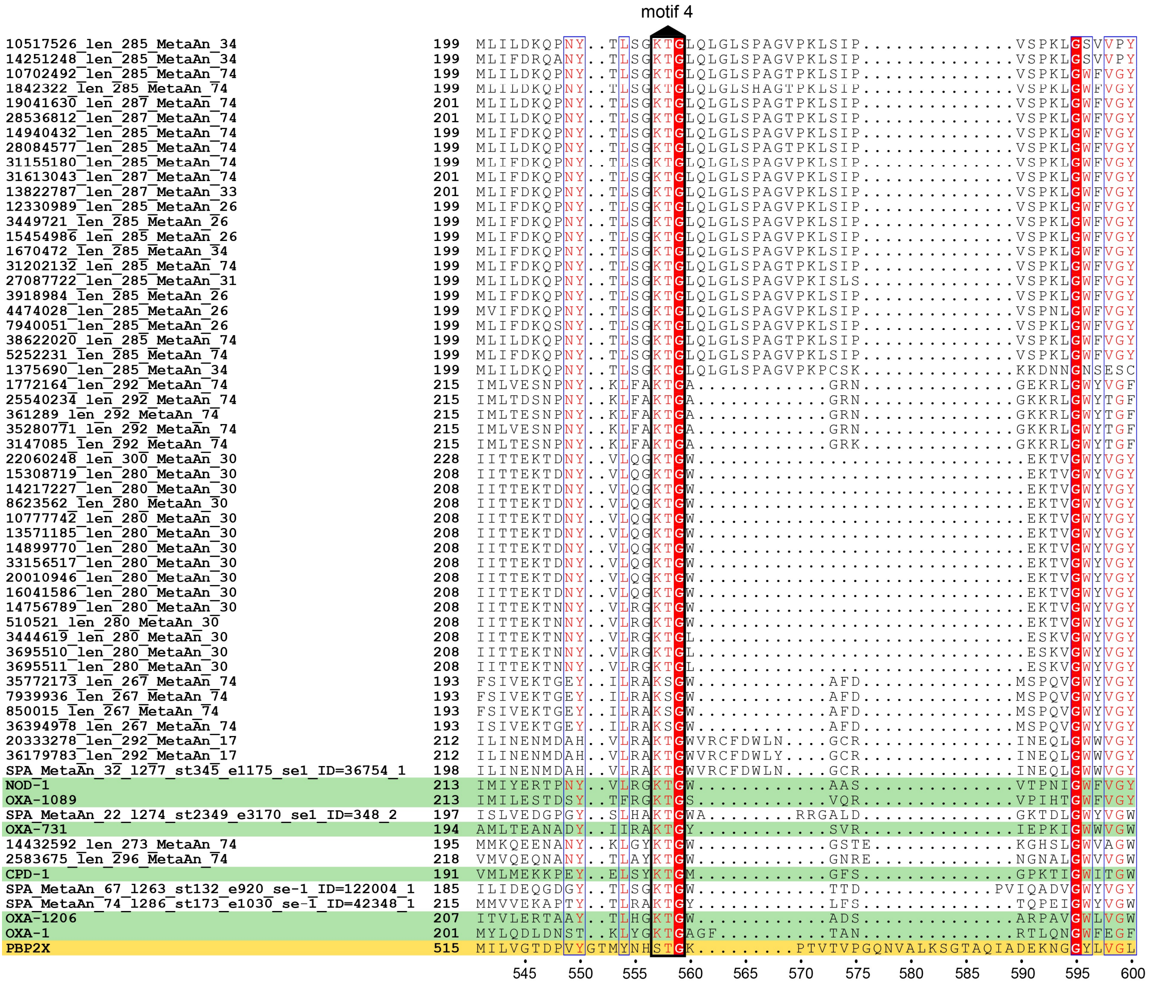


**Figure S3 (part 4/4).**

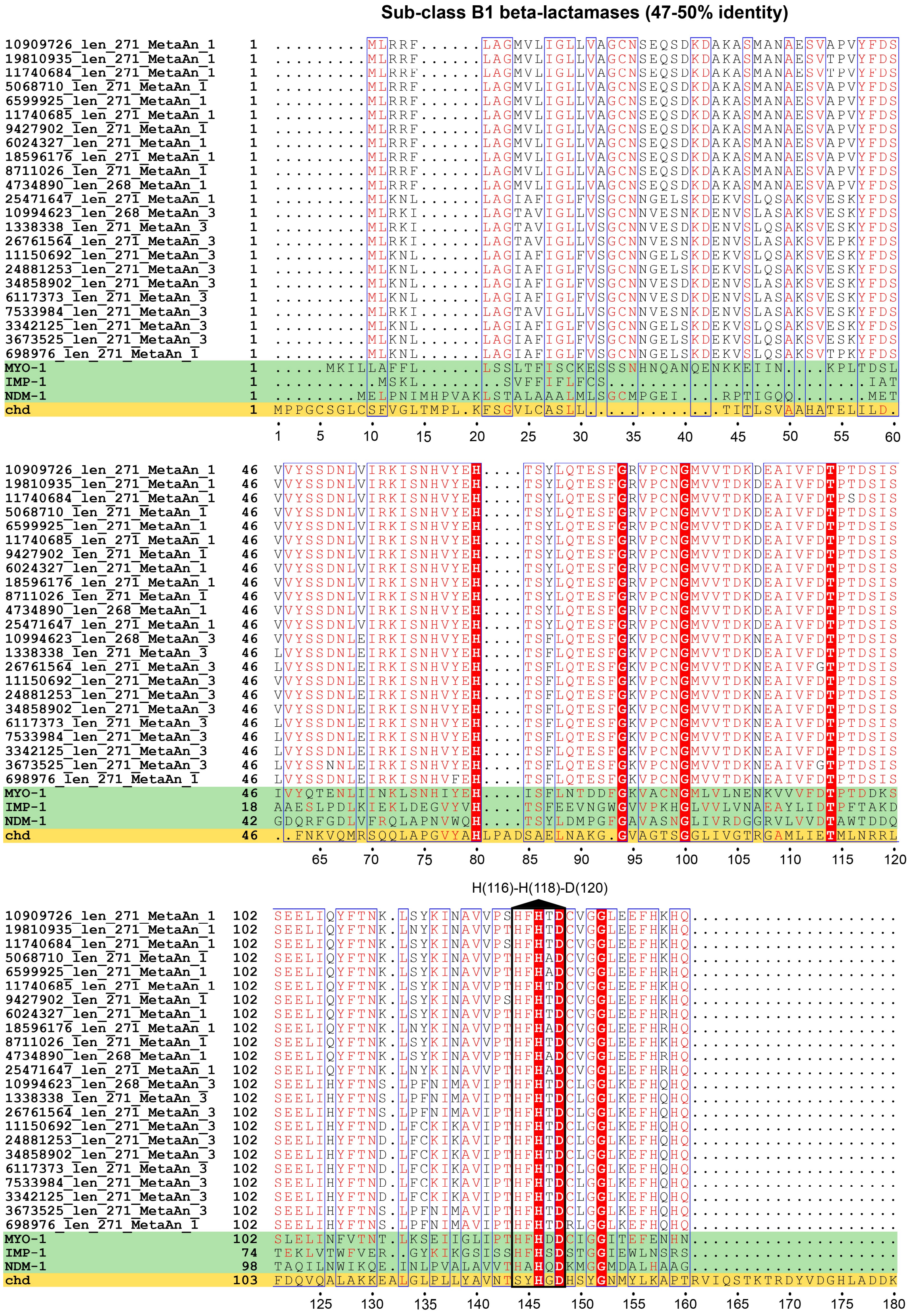


**Figure S4.** Multiple amino acid sequence alignment of key conserved regions among previously described class B1 beta-lactamases (shaded in green) and predicted Antarctic orthologs sharing 47-50% identity with the reference sequences. Chd protein (shaded in yellow) was included for comparison, as it belong to the MBL-like superfamily but does not have beta-lactamase activity (true negative).

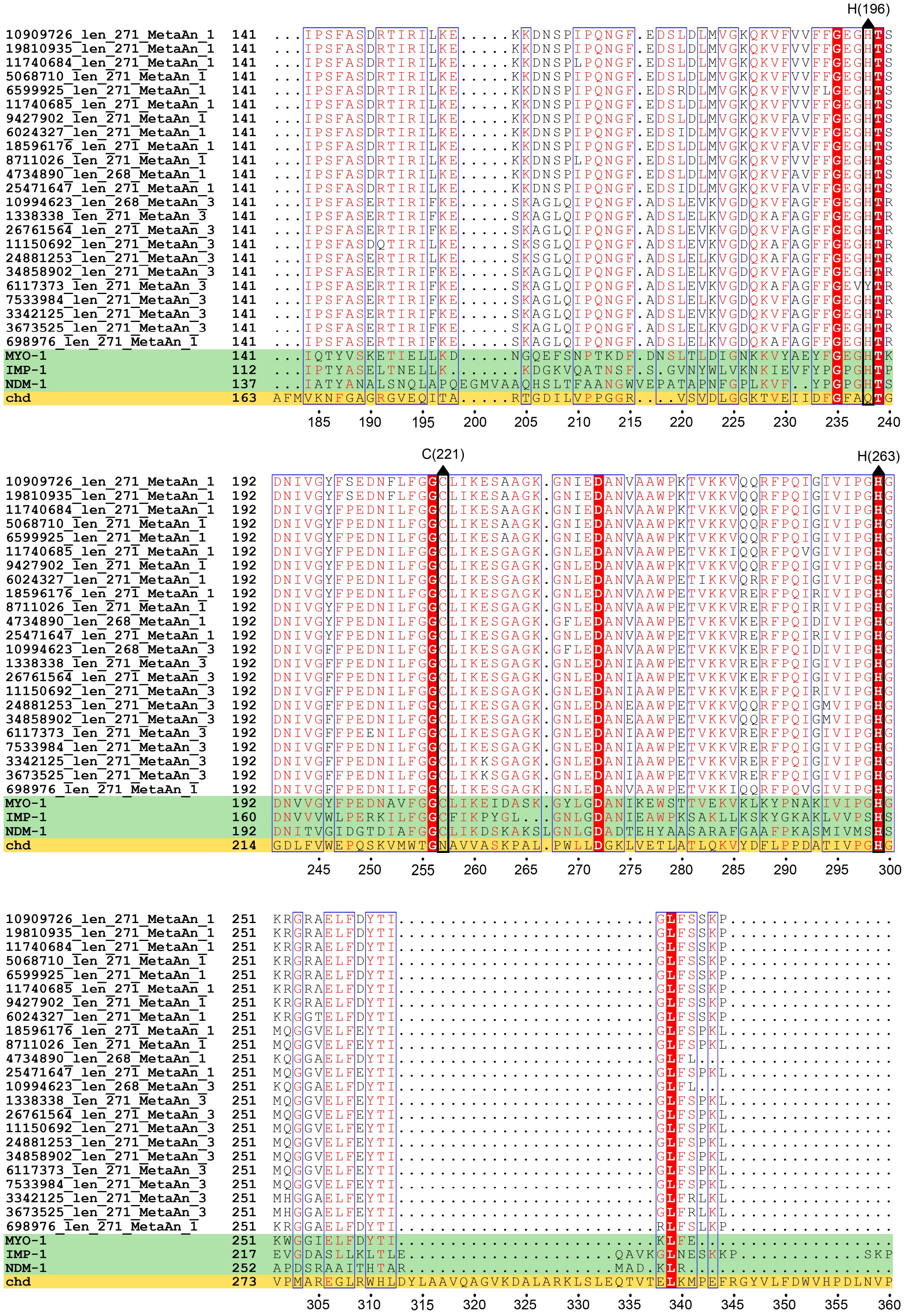


**Figure S4 (part 2/2).**

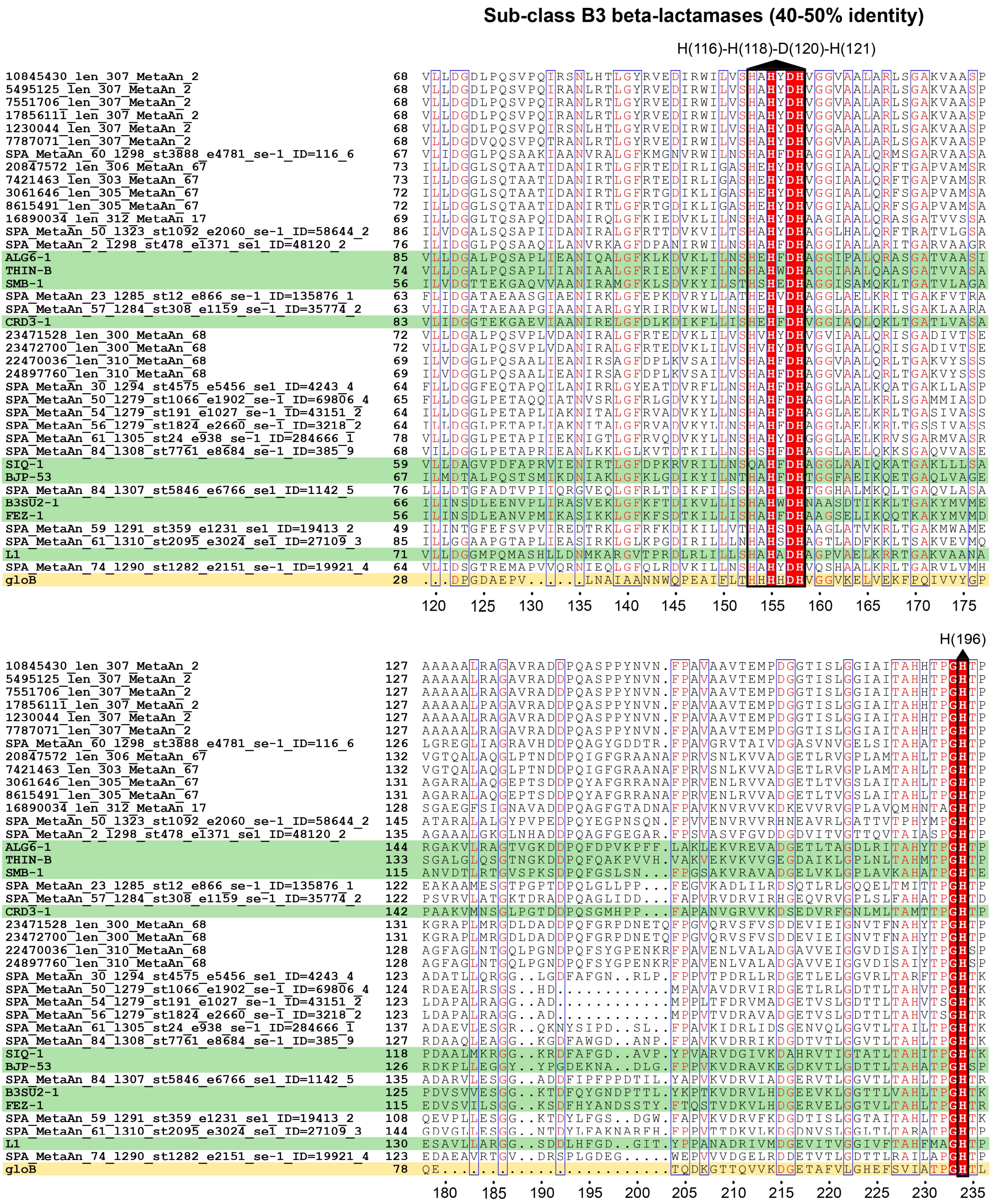


**Figure S5.** Multiple amino acid sequence alignment of key conserved regions among previously described class B3 beta-lactamases (shaded in green) and predicted Antarctic orthologs sharing 40-50% identity with the reference sequences. GloB protein (shaded in yellow) was included for comparison, as it belong to the MBL-like superfamily but does not have beta-lactamase activity (true negative).

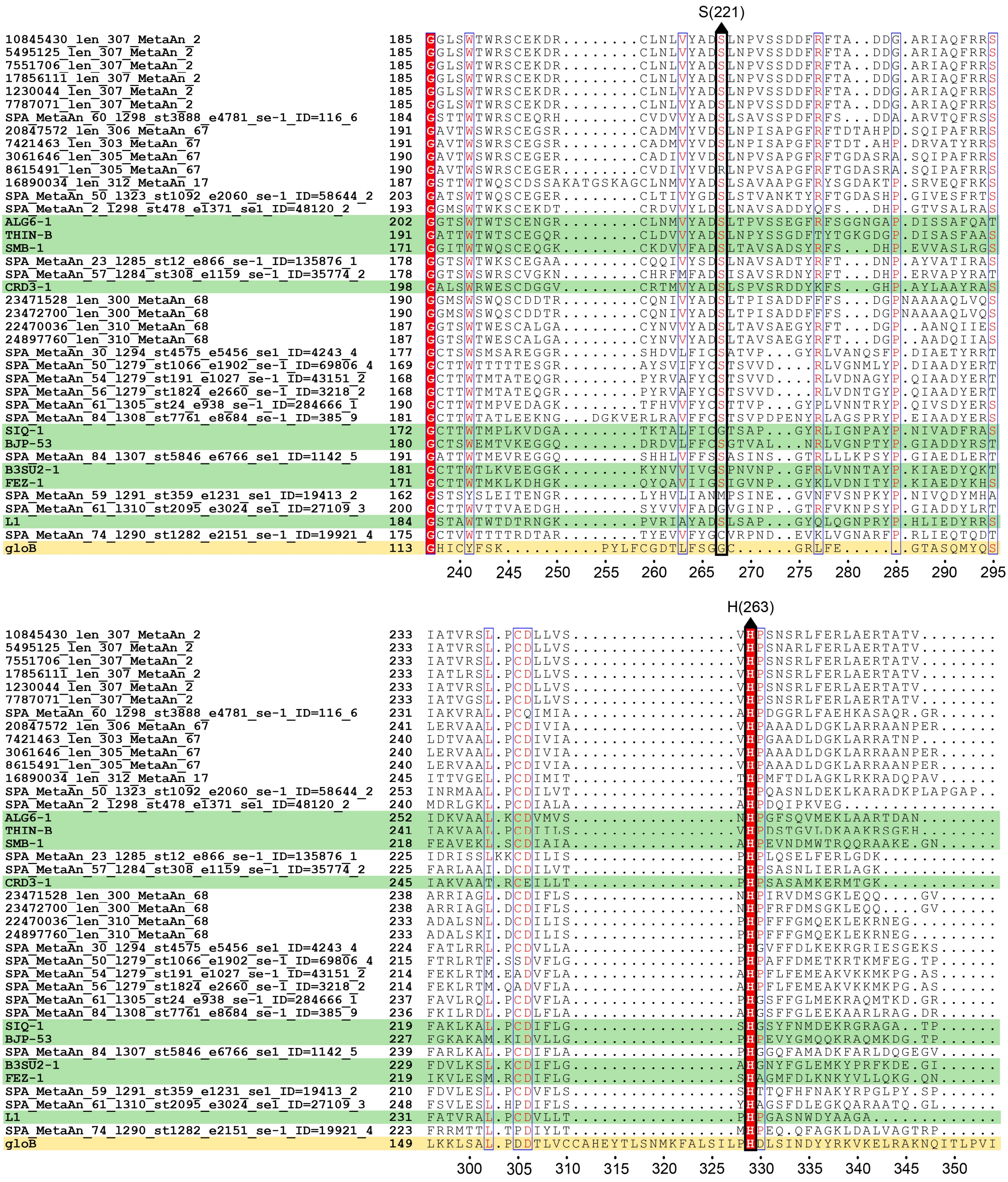


**Figure S5 (part 2/2).**

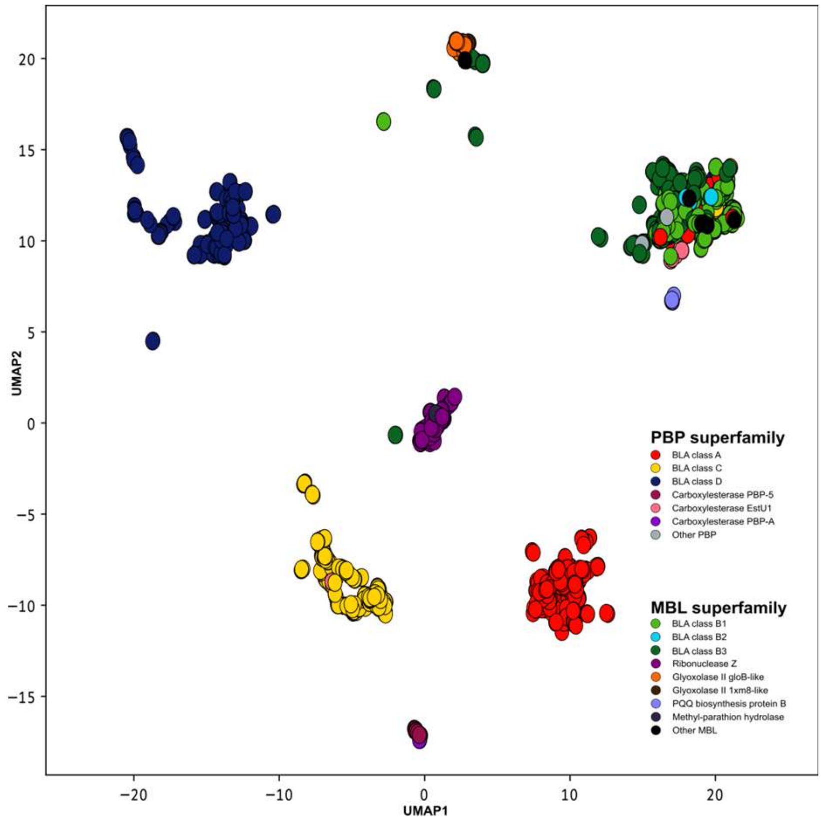


**Figure S6.** Uniform Manifold Approximation and Projection of functional similarities between references and Antarctic candidate BLAs and TNs. Each protein is represented as a 6109-dimensional GO-terms probability vector and cosines between vectors are used as metric for projection in two-dimensional space. Proteins are colored according to the observed protein class of reference proteins contained in each structural cluster.

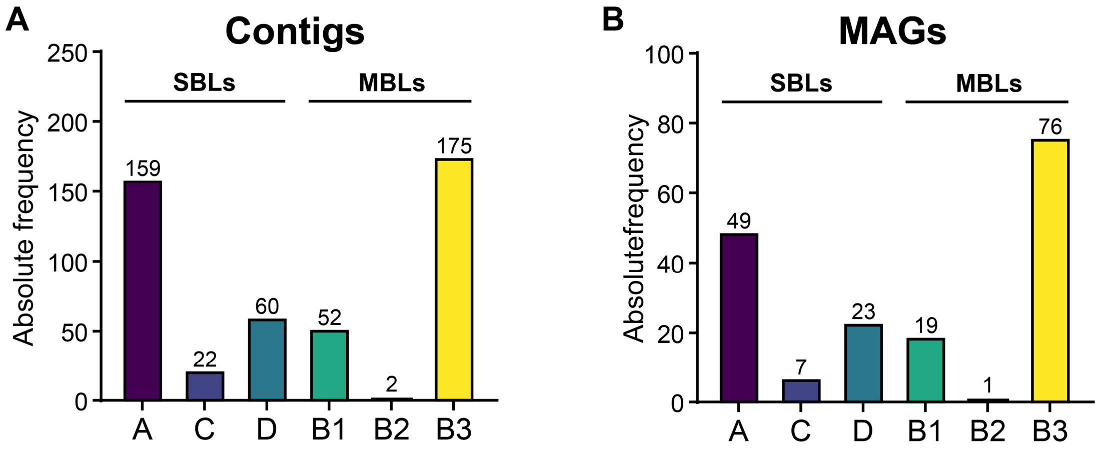


**Figure S7.** Beta-lactamase counts in assembled metagenomic contigs (A) and MAGs (B) from Antarctic and Subantarctic soils.

**Table S2.** PBP-like and MBL-like proteins lacking beta-lactamase activity used as negative control for Antarctic beta-lactamase orthologs search.

| Uniprot ID | Protein name | Annotation | Organism | Superfamily | Reference |
|---|---|---|---|---|---|
| P0AEB2 | PBP-5 | Carboxylesterase | *Escherichia coli* | PBP-like | Frohlich, 2021 |
| P72161 | PBP-A | Carboxylesterase | *Pseudomonas aeruginosa* | PBP-like | Frohlich, 2021 |
| K4HQE7 | EstU1 | Carboxylesterase | Metagenome | PBP-like | Frohlich, 2021 |
| P0AD70 | AmpH | Endopeptidase/carboxipeptidase | *Escherichia coli* | PBP-like | Frohlich, 2021 |
| P18357 | blaR1 | Transpeptidase | *Staphylococcus aureus* | PBP-like | Frohlich, 2021 |
| H8ZYG5 | PBP2X | Transpeptidase | *Streptococcus pneumoniae* | PBP-like | Frohlich, 2021 |
| Q988B9 | 3aj3 | 4-Pyridoxolactonase | *Mesorhizobium loti* | MBL-fold | Baier et al., 2014 |
| P0CJ63 | aiiA | AHL-lactonase | *Bacillus thuringiensis* | MBL-fold | Baier etal., 2014 |
| P28607 | atsA | Arylsulfatase | *Alteromonas carrageenovora* | MBL-fold | Baier et al., 2014 |
| C9EBR5 | chd | Chlorothalonil dehalogenase | *Pseudomonas aeruginosa* | MBL-fold | Baier etal., 2014 |
| Q9X466 | ncnE | Cyclase | *Streptomyces arenae* | MBL-fold | Daiyasu, 2001 |
| A0A096ZEC9 | dnhA | Dinitroanisole o-demethylase | *Nocardioides sp.* | MBL-fold | Fida etal., 2014 |
| Q50497 | fprA | Flavoprotein | *Methanothermobacter marburgensis* | MBL-fold | Daiyasu, 2001 |
| Q9SID3 | 1xm8 | Glyoxolase II | *Arabidopsis thaliana* | MBL-fold | Baier etal., 2014 |
| P0AC84 | gloB | Glyoxolase II | *Escherichia coli* | MBL-fold | Daiyasu, 2001 |
| Q9WZZ6 | 1ztc | Lactonase | *Thermotoga maritima* | MBL-fold | Baier etal., 2014 |
| Q841S6 | mpd | Methyl- parathion hydrolase | *Pseudomonas aeruginosa* | MBL-fold | Baier et al., 2014 |
| P16692 | phnP | Phosphate phosphodiesterase | *Escherichia coli* | MBL-fold | Daiyasu, 2001 |
| Q8DQ62 | lytD | Phosphorylcholine esterase | *Streptococcus pneumoniae* | MBL-fold | Baier et al., 2014 |
| Q88QV5 | pqqB | PQQ biosynthesis protein B | *Pseudomonas putida* | MBL-fold | Baier et al., 2014 |
| Q81U06 | 1zkp | Putative ribonuclease | *Bacillus anthracis* | MBL-fold | Baier et al., 2014 |
| P0A8V0 | rbn | Ribonuclease Z | *Escherichia coli* | MBL-fold | Baier etal., 2014 |
| Q9C8L4 | GLY3 | Sulfur dioxygenase | *Arabidopsis thaliana* | MBL-fold | Baier et al., 2014 |
| Q82ZZ3 | 2az4 | β-CASP ribonuclease | *Enterococcus faecalis* | MBL-fold | Baier etal., 2014 |

**Table S3.** Putative orthologs of true negatives from the PBP-like and MBL-fold-like superfamilies by protein class identified in 81 Antarctic metagenomes using the PROSSAF pipeline.

| Cluster name | Superfamily | Nº Antartic Sequences |
|---|---|---|
| Carboxylesterase EstU1 | PBP-like | 540 |
| Carboxylesterase PBP-5 | PBP-like | 138 |
| Carboxylesterase PBP-A | PBP-like | 8 |
| Glyoxolase II 1xm8- like | MBL-fold | 13 |
| Glyoxolase II gloB- like | MBL-fold | 161 |
| Methyl-parathion hydrolase | MBL-fold | 7 |
| PQQ biosynthesis protein B | MBL-fold | 33 |
| Pyridoxolactonase | MBL-fold | 4 |
| Ribonuclease Z | MBL-fold | 447 |
